# Chemical Boosting of Foldase Condensates Accelerates Oxidative Protein Folding

**DOI:** 10.64898/2026.07.08.737393

**Authors:** Mai Watabe, Tsubura Kuramochi, Momoka Fukushima, Misaki Kinoshita, Hiroki Akiba, Kazunori Ban, Mayuko Hashimoto, Noriyuki Uchida, Kenta Arai, Takakazu Nakabayashi, Johannes Buchner, Takahiro Muraoka, Masaki Okumura

## Abstract

Dynamic biomolecular condensates play crucial roles in intracellular compartmentalization and physiological functions. While engineering tools for compartmentalization have expanded add-on functionalities, directly amplifying the inherent catalytic machinery within biological phase-separated droplets has remained elusive. Herein, we developed a phase-separated oxidative folding reaction chamber based on protein disulfide isomerase A6 (PDIA6) by chemically targeting its active site CxxC motif to enhance enzymatic activity within PDIA6 droplets. A para-substituted N-methylated pyridinylmethanethiol (pMePySH) enhanced the catalytic oxidative folding of bovine pancreatic trypsin inhibitor, proinsulin, and antibody up to 12-fold within *in vitro* PDIA6 droplets. Furthermore, pMePySH targeted PDIA6 foci within the endoplasmic reticulum, significantly promoting insulin secretion. These findings offer a powerful platform for the spatiotemporal manipulation of protein folding, with profound implications for the scalable manufacturing of therapeutic antibodies and other complex biopharmaceuticals.

## Introduction

The spatiotemporal compartmentalization of chemical reactions within cells is crucial for separating and coordinating complex metabolic processes. Biomolecular condensates have emerged as key mediators of intracellular compartmentalization across various biological events, including synapsis formation ^1^, stress granules ^2^ ^3^, protein folding ^4,5^, protein degradation ^6^ ^7^, transport ^8^, and genome stability ^9^. By orchestrating specific biochemical pathways in a highly localized manner, biomolecular condensates offer a powerful blueprint for designing biomimetic reaction chambers for heterogeneous catalysis.

Nature has evolved diverse catalytic systems to control the formation of disulfide bonds in both eukaryotic and prokaryotic cells ^10^ ^11^. In eukaryotes, there are complex but sophisticated disulfide bond catalytic pathways within the endoplasmic reticulum (ER) that regulates disulfide bond formation-related protein folding ^12^ ^13^ ^14^. Recombinant expression of eukaryotic, disulfide-rich proteins and peptides such as proinsulin and antibodies in bacteria hosts like *Escherichia coli* often leads to the formation of inactive inclusion bodies due to aberrant disulfide bond pairing ^15–18^. Furthermore, the accumulation of proinsulin containing non-native disulfide pairings in the ER of human pancreatic cells is intimately linked to the pathogenesis of type 2 diabetes^19^. Therefore, dysregulation of the ER disulfide-catalytic machinery can lead to severe pathological outcomes^20^—a link that underscores the medical importance of understanding disulfide chemistry and highlights the promise of developing robust *in vitro* oxidative folding technologies for biopharmaceutical manufacturing.

The ER serves as a multi-functional organelle where protein folding, lipid synthesis, Ca^2+^ storage and post-translational modifications such as glycosylation and disulfide bond formation take place ^21^ ^22^ ^23^. It harbors more than 20 members of the protein disulfide isomerase (PDI) family, alongside key molecular chaperones such as immunoglobulin heavy-chain binding protein (BiP) and glucose-regulated protein 94 (GRP94), which collectively maintain the protein homeostasis network ^12^ ^24^ ^25^ ^26^. PDIA6, one member of the PDI family, is special because it forms liquid-like droplets in response to Ca^2+^; these condensates inhibit proinsulin aggregation and accelerate its oxidative folding, thereby promoting robust insulin secretion ^4^. These findings suggest a promising strategy wherein PDIA6 droplets can be leveraged as specialized reaction chambers for catalytic oxidative folding.

Despite considerable progress in developing efficient oxidative folding systems ^27^ ^28^ and expanding our biological understanding of the PDI family ^25^ ^26^ ^29^ (Fig. 1a,b), biotechnological concepts capable of fully controlling oxidative folding remain unrealized. To address this gap, we introduced a chemical biology strategy designed to target and accelerate enzymatic catalysis within PDIA6 droplets as potential reaction chambers for oxidative folding (Fig. 1c). Enhancing enzymatic activity directly within these protein quality control droplets is instrumental for accelerating the oxidative folding of therapeutic targets, such as insulin and antibodies. This approach combines the physical sequestration of unfolded substrates with creating a highly favorable microenvironment for disulfide bond formation (Fig. 1c). This work establishes a novel framework for manipulating protein folding and targeting protein quality control granules through the disulfide-catalyst function of PDIA6 condensates, providing a foundational strategy for boosting the activity of biological condensates using functionalized thiol compounds.

**Fig. 1.**
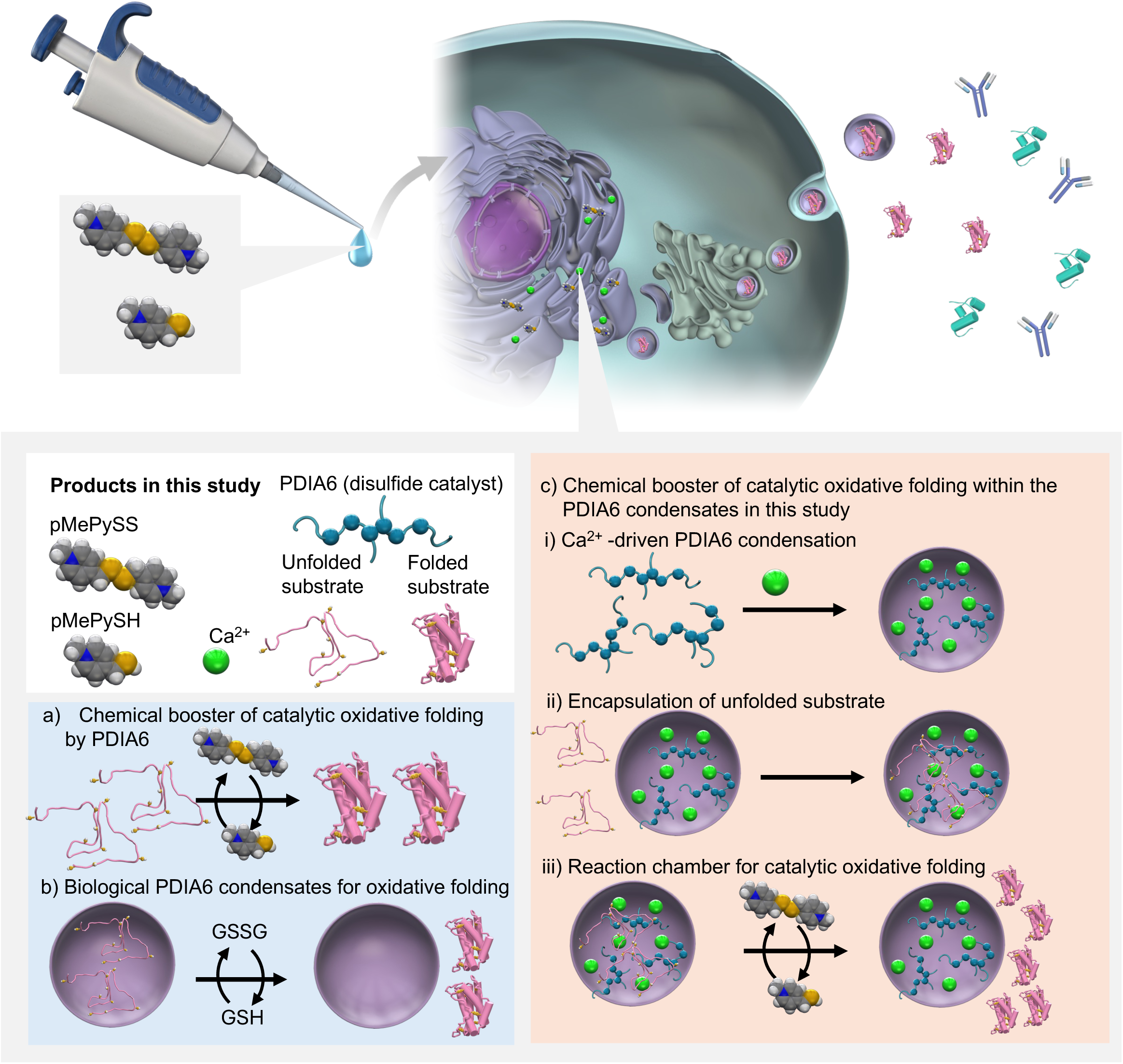
Concept behind this study. **a**, Chemical booster targeting of protein disulfide isomerase A6 (PDIA6), a disulfide catalyst, to enhance the catalysis of oxidative folding as reported previously. Para-substituted N-methylated pyridinylmethanethiol (pMePySH) enhances both the acidity and oxidizability of thiol groups, which allows dispersed PDIA6 to enhance its enzymatic activity in oxidative folding at a low loading compared to glutathione^35^. **b**, Liquid-like droplets serving as compartmentalization for protein quality control are formed by PDIA6 in a Ca^2+^-dependent manner to promote the correct folding of proinsulin while inhibiting its misfolding and aggregation^4^. **c**, Replacement of glutathione with the pMePySH redox system significantly enhances catalytic oxidative folding inside PDIA6 droplets, which serve as reaction chambers for the oxidative folding of substrates.

## Results

### PDIA6 droplets provide a general, enhanced disulfide-catalytic microenvironment for oxidative client folding

As a novel catalytic microenvironment for oxidative folding among the PDI family, PDIA6 biomolecular condensates have been shown to recruit reduced and denatured proinsulin, thereby both enhancing its oxidative proinsulin folding and inhibiting its aggregation ^4^. To investigate the generality of this enhanced catalytic microenvironment for other client proteins, we examined the oxidative protein folding of bovine pancreatic trypsin inhibitor (BPTI) ^30^ (Fig. 2a). BPTI is a well-established model protein containing three disulfide bonds, with a folding pathway that proceeds through well-defined intermediates (Fig. 2a). We prepared a BPTI mutant in which Cys5 was labeled with a DY-605-maleimide dye and all other cysteines were replaced with Ser. Because disulfide bonds stabilize local conformations to maintain tertiary structure, BPTI variants lacking these bonds adopt a random-coil conformation ^31^. This unfolded dye-labeled BPTI partitioned efficiently into PDIA6 droplets (Fig. 2b), mirroring the behavior of unfolded proinsulin ^4^.

**Fig. 2.**
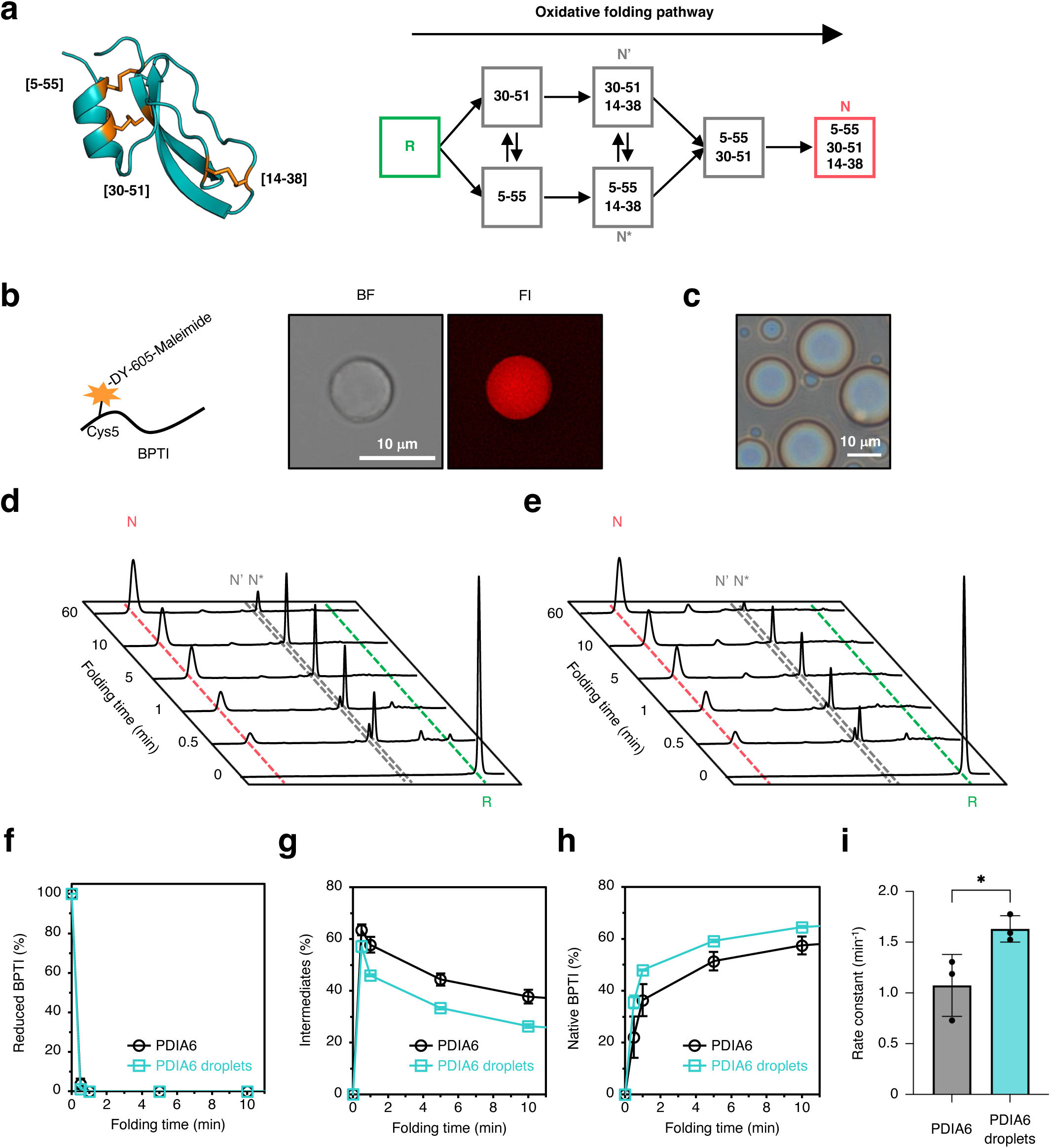
PDIA6 droplets as a reaction chamber for effective oxidative folding. **a**, Crystal structure of bovine pancreatic trypsin inhibitor (BPTI) (left). Disulfide bonds are shown as orange sticks. Schematic diagram of the BPTI oxidative folding pathway (right). R and N represent the reduced and native forms of BPTI (5–55, 14–38, and 30–51), respectively. Disulfide pairings of folding intermediates at each step are shown in boxes. **b**, Partitioning of dye-labeled BPTI into PDIA6 droplets. To prepare dye-labeled BPTI, Cys5 was labeled with DY-605-maleimide dye, and all other cysteines were replaced with Ser. Three independent experiments were performed. BF, bright-field image; FI, fluorescence image. This experiment was repeated three times independently. **c**, Microscopic imaging of PDIA6 droplets in the presence of reduced glutathione (GSH)/oxidized glutathione (GSSG). This experiment was repeated three times independently. **d,e**, Time-course reversed-phase high-performance liquid chromatography (HPLC) analysis of BPTI oxidative folding catalyzed by dispersed PDIA6 (**d**) and PDIA6 droplets (**e**). N, N’, N*, and R represent native BPTI, folding intermediates of BPTI with two disulfide pairings (30–51 and 14–38 for N’, and 5–55 and 14–38 for N*), and reduced forms of BPTI, respectively. Experiments were independently repeated three times. **f-h**, Time-course plots of reduced/denatured BPTI (**f**), folding intermediates of BPTI (**g**), and native BPTI (**h**) during BPTI folding by dispersed PDIA6 and PDIA6 droplets. These data were calculated from the HPLC areas of Fig. 2d,**e**. **i**, Folding kinetics of the formation rate of native BPTI in (**h**). Folding rate constants were determined from three independent experiments. Statistical significance was assessed using the two-tailed unpaired t-test. \**P*<0.05. Rate constants were calculated by single exponential curve fitting using IGOR Pro6 software (WaveMetrics, Inc., Lake Oswego, OR, USA, https://www.wavemetrics.com/) (Extended Data Fig. 1). **f-i**, Data are presented as the mean ± s.d.

We next evaluated the kinetics of BPTI oxidative folding within PDIA6 droplets. Native BPTI contains six Cys residues (Cys5, Cys14, Cys30, Cys38, Cys51, and Cys55) that form three native disulfide bond pairings: Cys5–Cys55, Cys14–Cys38, and Cys30–Cys51 (Fig. 2a). The oxidative folding of BPTI from its reduced state to the native conformation involves kinetically trapped several folding intermediates such as N’, N*, and N ^SH^ (Fig. 2a), which can be resolved and monitored via reversed-phase high-performance liquid chromatography (RP-HPLC) ^30^. Reduced and denatured BPTI (5 µM) was incubated with PDIA6 (20 µM) —a concentration well within the regime for Ca^2+^-induced liquid-like droplet formation ^4^—under standard redox conditions ([GSH] = 2 mM and [GSSG] = 0.1 mM) in the presence or absence of Ca^2+^ (Fig. 2c–e).

RP-HPLC analysis revealed that whereas in the presence of dispersed PDIA6 ∼20% native BPTI were generated within 30 s, PDIA6 droplets yielded ∼40% native BPTI over the same period. After 60 min, the folding reaction approached completion within PDIA6 droplets, whereas intermediates continued to accumulate in the presence of dispersed PDIA6 (Fig. 2d,e). Kinetic profiling of the disappearance of reduced BPTI and the accumulation of intermediates and native species yielded apparent rate constants of *k*_dispersed_PDIA6_ = 1.0 min^-1^ and *k*_PDIA6_condensates_ = 1.6 min^-1^ (Fig. 2f–i and Extended Data Fig. 1). This demonstrates that phase separation enhances the catalytic efficiency of PDIA6 towards BPTI by approximately 1.6-fold. This acceleration is consistent with our previous finding that PDIA6 condensation enhances catalytic activity toward proinsulin by ∼2.6-fold ^4^. Collectively, these results underscore the potential of leveraging PDIA6 condensates as specialized reaction chambers for encapsulation and oxidative folding of unstructured client proteins.

### Substrate partitioning into PDIA6 foldase condensates depends on folding states

To further dissect the enzymatic mechanisms operating within PDIA6 foldase condensates, we systematically evaluated how substrate concentration and folding states influence droplet properties. Three-dimensional refractive index (RI) tomography allows for the non-invasive visualization of the internal macromolecular packing density within PDIA6 droplets over time ^4^. Experiments were performed below the critical concentration for autonomous PDIA6 condensation (50 µM) by utilizing Ca^2+^ (4 mM) ^4^ to exclude the effect of substrate binding by dispersed PDIA6. In the absence of client proteins, the internal RI values and fusogenicity of PDIA6 condensates remained stable over 60 min (Fig. 3a). Upon the addition of native BPTI, the internal RI of the condensates increased significantly in a dose-dependent manner, demonstrating robust and selective encapsulation of the folded substrate (Fig. 3b,c).

**Fig. 3.**
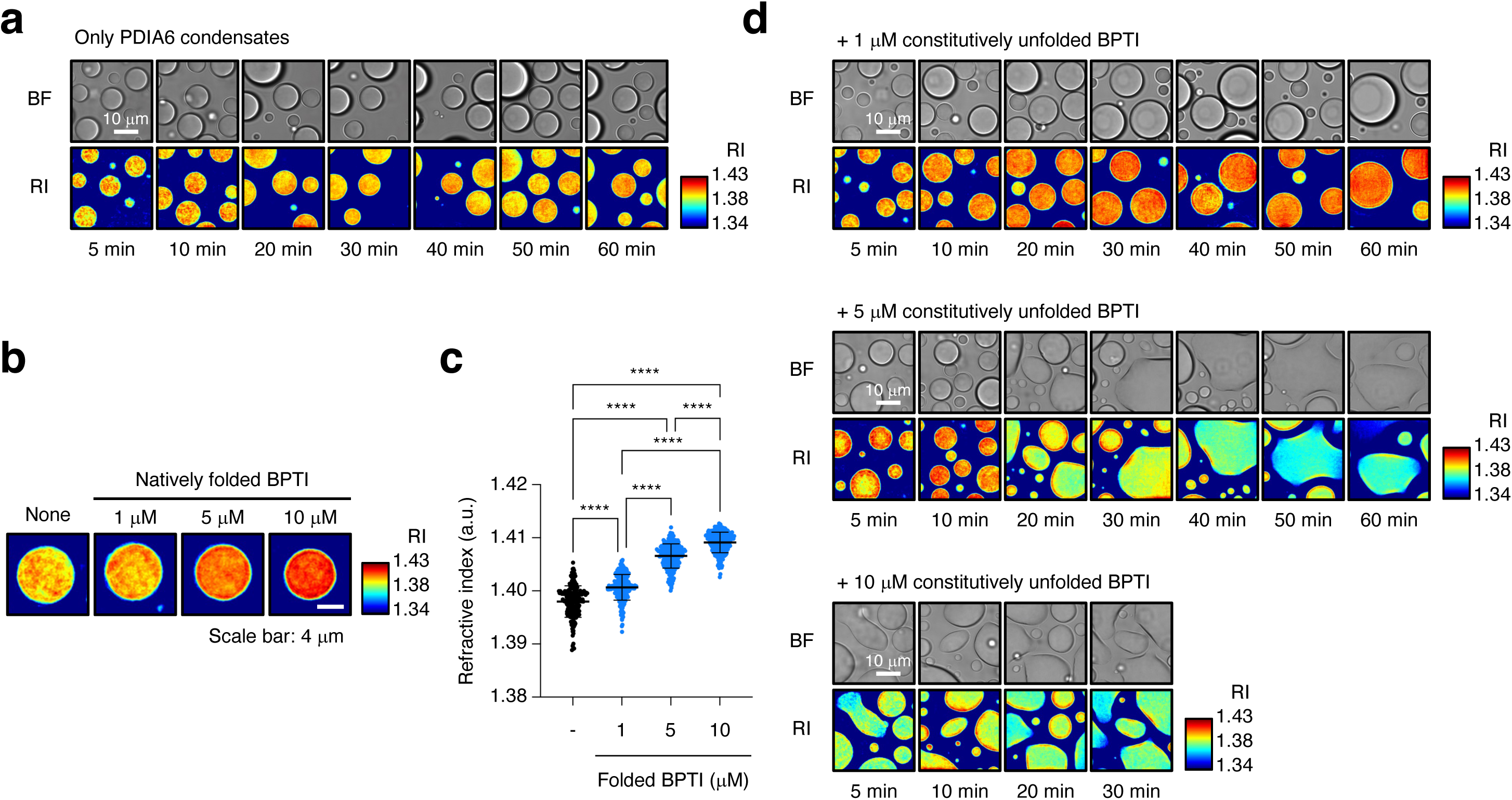
Uptake capability into PDIA6 condensates for native and unfolded BPTI. **a**, Representative bright-field (top) and two-dimensional (2D) refractive index (RI) distribution images (bottom) of PDIA6 condensates over time. Experiments were independently repeated three times. **b,** Representative 2D RI distribution images within PDIA6 condensates in the absence or presence of 1–10 µM native BPTI. Three independent experiments were performed. **c**, Statistical analyses of RI within PDIA6 condensates in the absence or presence of native BPTI. Experiments were independently repeated at least three times (n = 206 particles for only PDIA6 condensates, 239 particles for 1 µM BPTI, 208 particles for 5 µM BPTI and 246 particles for 10 µM BPTI). Data are presented as the mean ± s.d. Statistical significance was examined using a one-way ANOVA with Tukey’s honest significant difference post-hoc test; the test was two-sided. \*\*\*\**P*<0.0001. **d**, Representative bright-field and 2D RI distribution images of PDIA6 condensates with 1–10 µM constitutively unfolded BPTI over time. This experiment was replicated three times independently.

With the goal of developing a targeting strategy to catalyze the folding of denatured substrates within these condensates, we used the constitutively unfolded random-coil control using a BPTI mutant in which all cysteine residues were substituted with serine ^31^. Time-course RI profiling revealed that the droplet interiors became progressively denser when incubated with either folded BPTI or a low concentration (1 μM) of constitutively unfolded BPTI (Fig. 3a,d and Extended Data Fig. 2a). In contrast, exposure to 5 µM of constitutively unfolded BPTI initially increased internal packing up to 10 min, after which the macromolecular assembly was disrupted. Furthermore, higher concentrations (10 µM) of the unfolded mutant triggered a rapid collapse of the PDIA6 condensates (Fig. 3d).

While many biomolecular condensates are driven by weak, multivalent interactions between low-complexity domains (LCDs) ^32,33^, we previously established that PDIA6 phase separation relies on a transient, specific, and electrostatic multivalent network between its first (redox-active) and third (redox-inactive) thioredoxin (Trx)-like domains ^4^. Our current findings suggest a molecular mechanism wherein an excess of random-coil structures—resembling intrinsically disordered regions (IDRs)—interferes with these critical inter-domain electrostatic interactions, thereby destabilizing the condensate architecture. Importantly, unlike the constitutively unfolded mutant, the addition of an equivalent concentration of foldable, reduced/denatured BPTI did not cause condensate collapse, even after 30 min of incubation (Extended Data Fig. 2b). This disparity implies that active catalytic turnover within the condensates continuously depletes the pool of unfolded substrates, thereby preserving the internal interaction network and maintaining the reversible material properties of the condensates. These findings emphasize that maintaining high catalytic turnover within PDIA6-based reaction chambers is a prerequisite for preserving condensate integrity, preventing client-induced structural dissolution during bioprocess operations.

### Addition of pMePySH/pMePySS hyper-activates catalytic oxidative BPTI folding within PDIA6 droplets

In the ER, glutathione drives thiol-disulfide exchange reactions, where both its reduced (GSH) and oxidized (GSSG) forms regulate the redox status of PDI family oxidoreductases, suggesting that small-molecule thiols can serve as potential switches. We previously demonstrated that para-substituted N-methylated pyridinyl-methanethiol (pMePySH; p*K*a = 7.34, E°’ = -211 mV) ^34^, which possesses higher thiol acidity and a more oxidizing redox potential than glutathione (p*K*a = 9.17, E°’ = -256 mV), substantially enhances the catalytic efficiency of dispersed PDIA6 ^35^. This prominent activation at low molar ratios prompted us to explore whether enzymatic activity within PDIA6 droplets—which provide a highly concentrated catalytic microenvironment—could be similarly manipulated via their active-site CxxC motif. We first evaluated the impact of pMePySH/pMePySS on the formation of PDIA6 condensation and observed liquid-like droplet behavior comparable to that seen with glutathione (Fig. 4b,c). This confirmation indicated that pMePySH/pMePySS does not interfere with the structural integrity of the condensate reaction chambers *in vitro*.

**Fig. 4.**
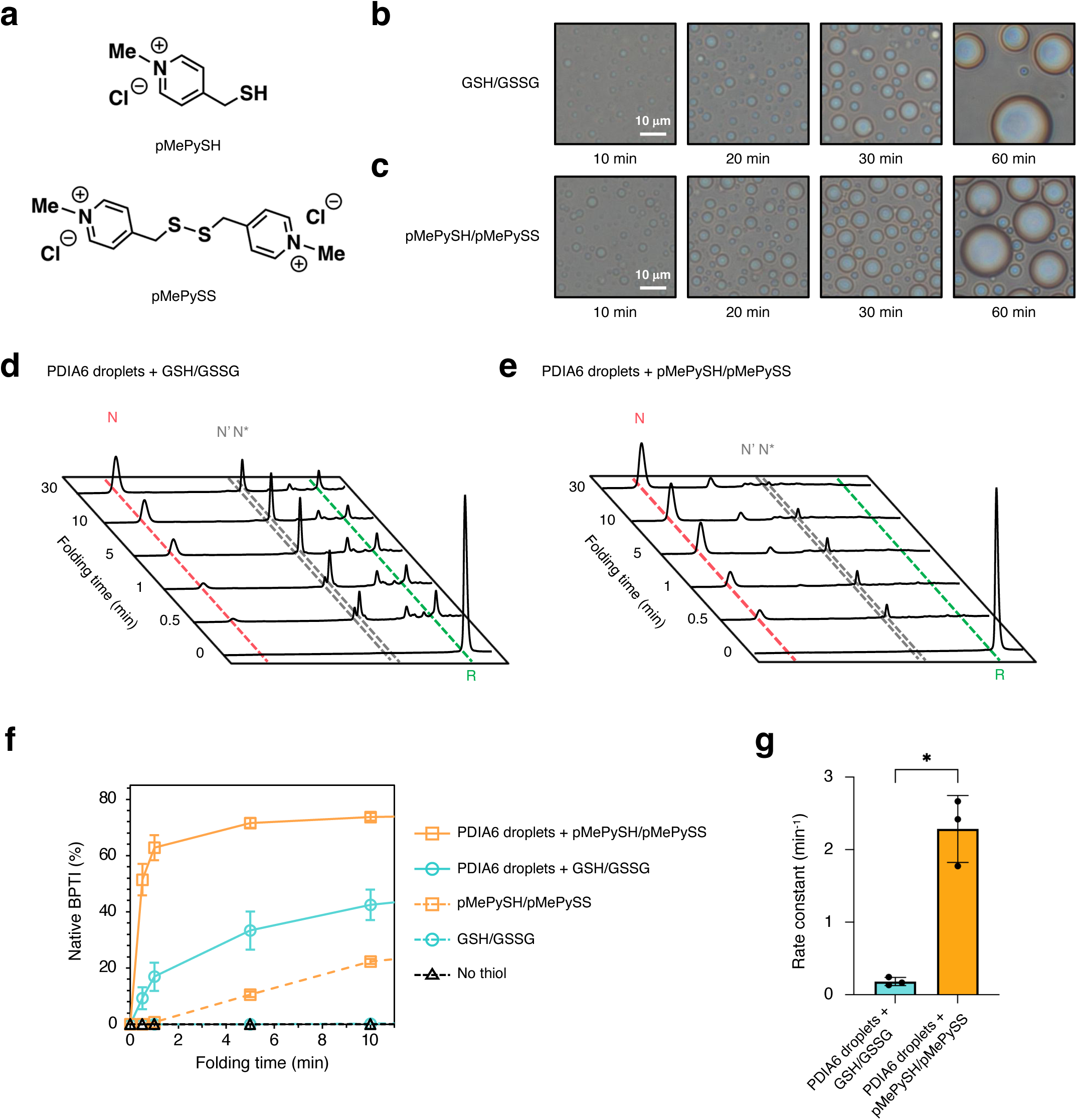
Catalytic oxidative BPTI folding inside PDIA6 droplets is enhanced 12-fold by pMePySH/pMePySS compared with GSH/GSSG. **a**, Chemical structures of pMePySH (top) and pMePySS (bottom). **b,c**, Time-course microscopic images of PDIA6 droplet formation in the presence of GSH/GSSG (**b**) or pMePySH/pMePySS (**c**). These experiments were replicated three times independently. **d,e**, Time-course reversed-phase HPLC analysis of oxidative BPTI folding by PDIA6 droplets in the presence of GSH/GSSG (**d**) or pMePySH/pMePySS (**e**). N, N’, N*, and R represent native BPTI, folding intermediates of BPTI with two disulfide pairings (30–51 and 14–38 for N’, and 5–55 and 14–38 for N*), and reduced/denatured forms of BPTI, respectively. **d,e**, These experiments were performed three times independently. **f**, Time-course plots of the formation of native BPTI (N) during BPTI folding catalyzed by GSH/GSSG in (**Extended Data Fig. 3a**), pMePySH/pMePySS in (**Extended Data Fig. 3b**), PDIA6 droplets in (**Extended Data Fig. 3c**), PDIA6 droplets with GSH/GSSG in (**d**) or pMePySH/pMePySS in (**e**). **g**, Folding kinetics of the formation rate of native BPTI in (**f**). Rate constants were calculated by single exponential curve fitting using IGOR Pro6 software (Extended Data Fig.3e,f). Statistical significance was assessed using the two-tailed unpaired t-test. \**P*<0.05. **f,g**, Data are presented as the mean ± s.d.

We next monitored the oxidative folding of BPTI inside PDIA6 droplets in the presence of these redox couples ([BPTI] = 5 μM, [disulfides] = 5 μM, [thiols] = 75 μM). To evaluate the catalytic turnover driven by PDIA6 condensates, we utilized pMePySS as the oxidant at a concentration lower than the total thiol content of reduced BPTI. In the absence of PDIA6 droplets, pMePySH/pMePySS accelerated native BPTI formation more effectively than GSH/GSSG within 5 min (Extended Data Fig. 3a,b), confirming the efficiency of this low substrate-to-compound ratio ^35^. Crucially, while PDIA6 droplets alone lacked the capacity to initiate disulfide bond formation (Extended Data Fig. 3c), the combination of PDIA6 droplets and pMePySH/pMePySS triggered the rapid and near-complete disappearance of reduced BPTI within 30 s. In contrast, ∼20% of BPTI remained in the reduced form under the GSH/GSSG droplet condition (Fig. 4d, e). Simultaneously, the pMePySH/pMePySS-supplemented droplets yielded a 6-fold increase in native BPTI production after 30 s mark compared to the GSH/GSSG droplets (Fig. 4d–f and Extended Data Fig. 3d). Notably, while independent incubation with pMePySH/pMePySS alone yielded only ∼20% native BPTI after 10 min (Fig. 4f and Extended Data Fig. 3b, d), its synergy with PDIA6 droplets drove native BPTI production to ∼80% over the same period (Fig. 4e, f and Extended Data Fig. 3d). Kinetic plots of native BPTI formation yielded apparent rate constants of *k*_condensates_GSH/GSSG_ = 0.2 min^-1^ and *k*_condensates_pMePySH/pMePySS_ = 2.3 min^-1^ (Fig. 4g), demonstrating that pMePySH/pMePySS hyper-activates the enzymatic machinery within PDIA6 droplets by approximately 12-fold. Leveraging its superior thiol acidity and optimized redox potential relative to glutathione, pMePySH effectively primes the highly concentrated enzymatic microenvironment within PDIA6 droplets for enhanced catalytic turnover.

### Oxidative folding of other client proteins within foldase condensates is synergistically enhanced by pMePySH/pMePySS

To establish the generality of this enhanced catalytic platform, we investigated the synergistic effects of PDIA6 droplets and pMePySH/pMePySS on the oxidative folding of diverse client proteins, including D3-L11^36^ (anti-hen egg lysozyme (HEL) VHH domain) and human proinsulin—a therapeutic target of high biopharmaceutical relevance. To evaluate oxidative VHH folding within the reaction chamber, we monitored reduced and denatured D3-L11, which requires the formation of a single disulfide bond (Fig. 5a and Extended Data Fig. 4a,b) ^37^ . The pMePySH/pMePySS-potentiated chambers produced ∼75% native D3-L11 within 30 s, whereas PDIA6 droplets under GSH/GSSG conditions yielded only ∼40% native species at the same time point (Fig. 5b and Extended Data Fig. 4c,d).

**Fig. 5.**
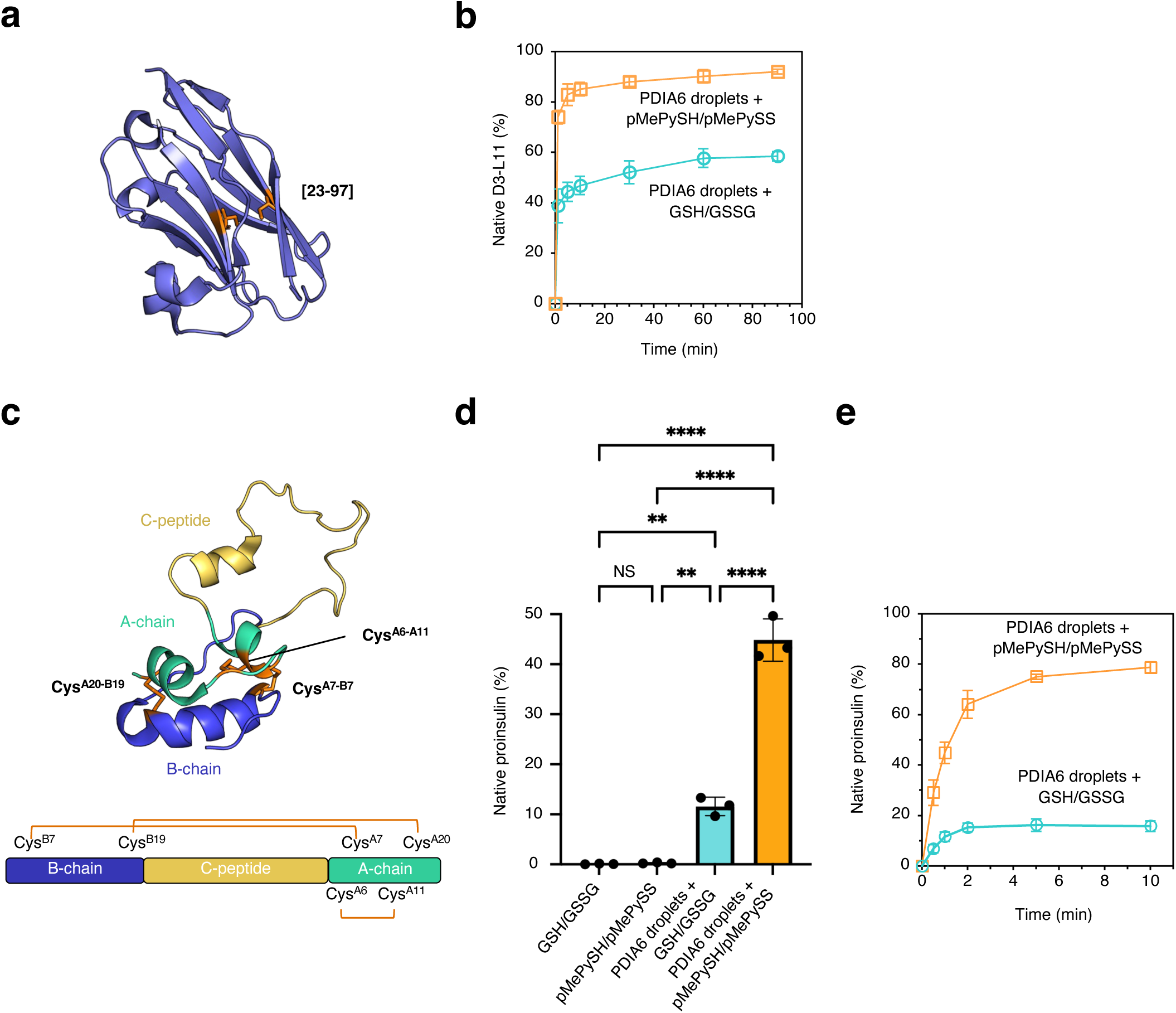
Synergistically enhanced oxidative folding inside the foldase condensates by PDIA6 in combination with pMePySH/pMePySS. **a**, Crystal structure of D3-L11 (PDB code: 6JB9), which contains a single disulfide bond. **b**, Time-course plot of native D3-L11 formation catalyzed by PDIA6 droplets in the presence of GSH/GSSH or pMePySH/pMePySS. **c**, Solution NMR structure of proinsulin (PDB code: 2KQP). Native proinsulin contains three disulfide bonds (CysB7–CysA7, CysB19–CysA20, and CysA6–CysA11). **d**, Oxidative folding yields of native proinsulin after 1 min in the presence of GSH/GSSG or pMePySH/pMePySS, with or without PDIA6 condensates. Statistical significance was examined with ANOVA with Tukey’s honest significant difference post-hoc test; the test was two-side. \*\**p*<0.01; \*\*\*\**p*<0.0001; NS, not significant. **e**, Time-course plot of native proinsulin formation for catalysis by PDIA6 droplets in the presence of GSH/GSSH or pMePySH/pMePySS. **b,d,e**, Data are presented as the mean ± s.d of three independent experiments.

Under physiological conditions, nascent human proinsulin is translocated into the ER where two interchain disulfide bonds (CysB7-CysA7 and CysB19-CysA20) and one intrachain disulfide bond (CysA6-CysA11) are introduced to stabilize its tertiary conformation (Fig. 5c)^38^ ^39^. Although *in vitro* proinsulin folding is highly inefficient at the physiological ER pH of 7.2 ^35^, our recent work demonstrated that PDIA6 droplets function as ER-localized disulfide catalysts that encapsulate proinsulin and accelerate its oxidative folding in a GSH/GSSG-dependent manner, a process vital for insulin secretion ^4^. Indeed, PDIA6 droplets promoted native proinsulin formation even at lower GSH/GSSG concentrations than previously reported (Fig. 5d) ^4^. Consistent with the BPTI results, PDIA6 droplets lacking redox molecules failed to initiate disulfide formation in proinsulin (Extended Data Fig. 4e). However, the addition of pMePySH/pMePySS to PDIA6 droplets significantly outpaced the GSH/GSSG condition in generating native proinsulin (Fig. 5d,e). The observed folding acceleration markedly exceeded the cumulative effects of the independent components, confirming a robust synergistic mechanism for proinsulin folding within the condensate microenvironment.

### pMePySH/pMePySS targets PDIA6 foci in U2OS cells to promote insulin secretion

Having established that pMePySH/pMePySS significantly promotes the *in vitro* oxidative folding of diverse proteins (Figs. 4f, 5b,e), we next evaluated the effects of pMePySH in cells. To clarify drug penetration into cells, we synthesized a fluorescently labeled pMePySH derivative tagged with bromobimane (pMePySH-BBr) (Fig. 6a and Extended Data Fig. 5a). Notably, when added to cells, pMePySH-BBr co-localized with mCherry-PDIA6 foci within the ER, confirming that the chemical booster successfully targets PDIA6 condensates *in cellulo* (Fig. 6b and Extended Data Fig. 5b). Treatment of U2OS cells stably expressing mCherry-PDIA6 with 75 μM pMePySH/5 μM pMePySS for 8 h exerted negligible effects on the morphology or number of intracellular PDIA6 foci relative to the GSH/GSSG control (Fig. 6c–e). While excessive oxidative stress—such as exposure to 5 mM hydrogen peroxide (H_2_O_2_)—triggered apoptosis via caspase-3/7 activation, pMePySH/pMePySS induced no cytotoxicity at the concentrations used (75 μM of reducing agent and 5 μM of oxidizing agent), demonstrating excellent biocompatibility (Fig. 6d, e).

**Fig. 6.**
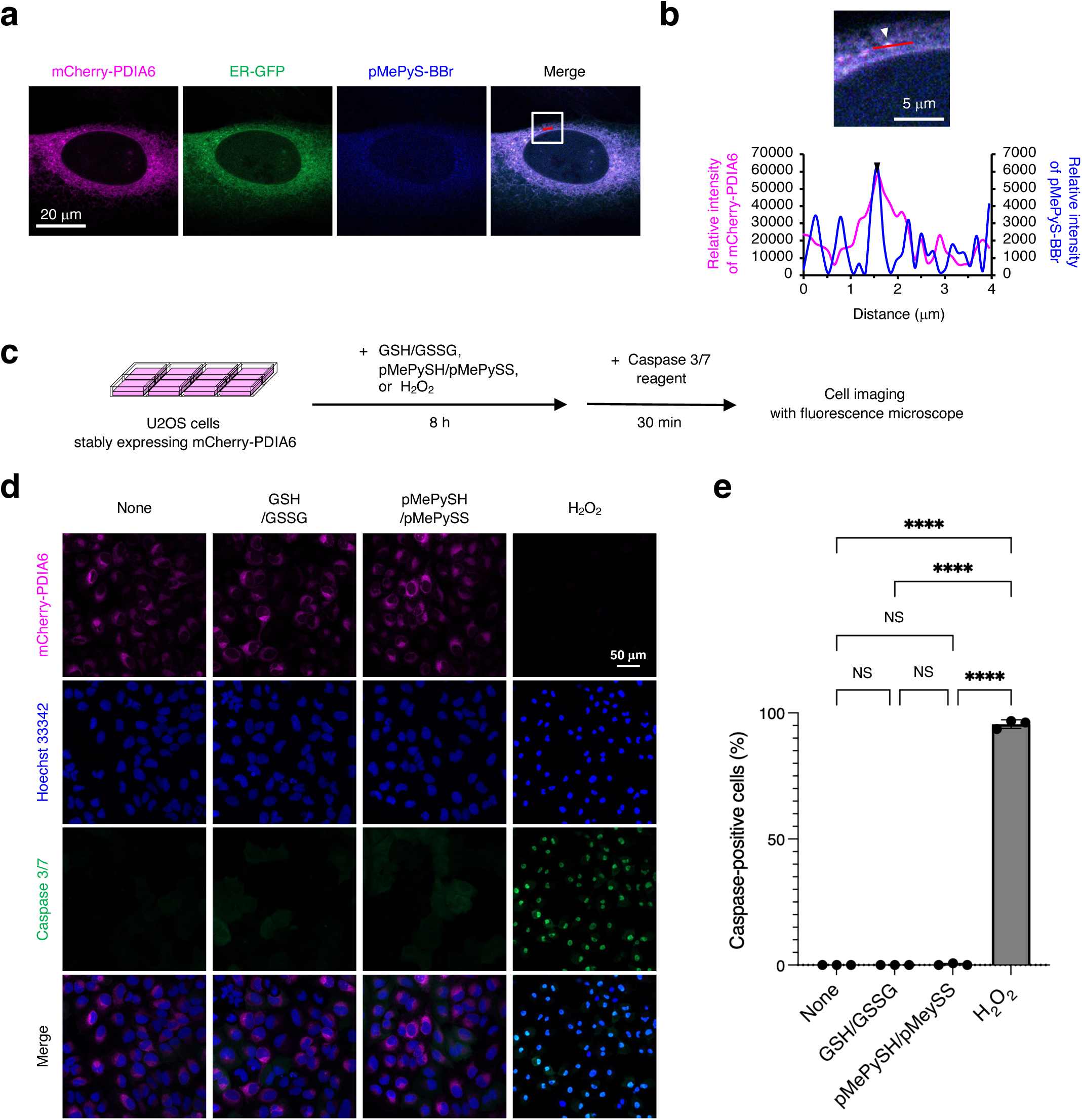
pMePySH co-localizes with PDIA6 in U2OS cells. a, Co-localization of stably expressed PDIA6 and pMePySH. U2OS cells stably expressing mCherry-PDIA6 were stained with CellLight ER-GFP and were treated with 75 µM fluorescently labelled pMePySH (pMePyS-BBr) for 5 h. **b**, Magnified views of the regions demarcated by white squares in **a** (top). The white arrowhead indicates specific foci selected for intensity profiling. Line scan analysis showing the relative fluorescence intensities of mCherry-PDIA6 (magenta) and pMePyS-BBr (blue) along the red line indicated in the magnified image (bottom). The black arrowhead in the line profile corresponds to the position of the PDIA6 focus indicated by the white arrowhead in the top panel. **c**, Schematic of foci formation and the cellular apoptosis assay used to evaluate the effects of GSH/GSSG, pMePySH/pMePySS, and H2O2 on U2OS cells stably expressing mCherry-PDIA6. **d,e**, Representative images (**d**) and quantification (**e**) of U2OS cells stably expressing mCherry-PDIA6 treated with or without 75 µM GSH/ 5 µM GSSG, 75 µM pMePySH/5 µM pMePySS and 5 mM H_2_O_2_ for 8 h. The caspase-positive cells were detected with CellEvent Caspase-3/7 Detection Reagents. Nuclei were stained with Hoechst 33342. **e**, Data are presented as the mean ± s.d. Statistical significance was examined using a one-way ANOVA with Tukey’s honest significant difference post-hoc test; the test was two-sided. \*\*\*\**P*<0.0001; NS, not significant. **a,b,d,e,** These experiments were performed three times independently.

To quantify the physiological impact of this system on the secretion of exogenously expressed proinsulin, we generated *PDIA6*-knockout (KO) HCT116 cells. Consistent with our previous findings ^4^, *PDIA6*-KO cells transfected with a proinsulin-Gaussia luciferase fusion (Proinsulin-Gluc) exhibited significantly reduced insulin secretion, which was fully rescued upon the reintroduction of PDIA6 (Fig. 7a,b). This confirms that ER-localized PDIA6 condensates serve a critical quality control function essential for the oxidative folding and export of insulin. Remarkably, while the addition of pMePySH/pMePySS to *PDIA6*-KO cells did not alter baseline insulin secretion, its administration to PDIA6-rescued cells significantly enhanced insulin secretion (Fig. 7b). Importantly, the secretory deficit inherent to PDIA6-KO cells was not mitigated by pMePySH/pMePySS administration alone; rather, the chemical booster required the physical presence of PDIA6 condensates to augment insulin export. Taken together, these data demonstrate that pMePySH/pMePySS successfully targets foldase condensates within the ER to enhance the efficiency of oxidative protein folding *in cellulo*.

**Fig. 7.**
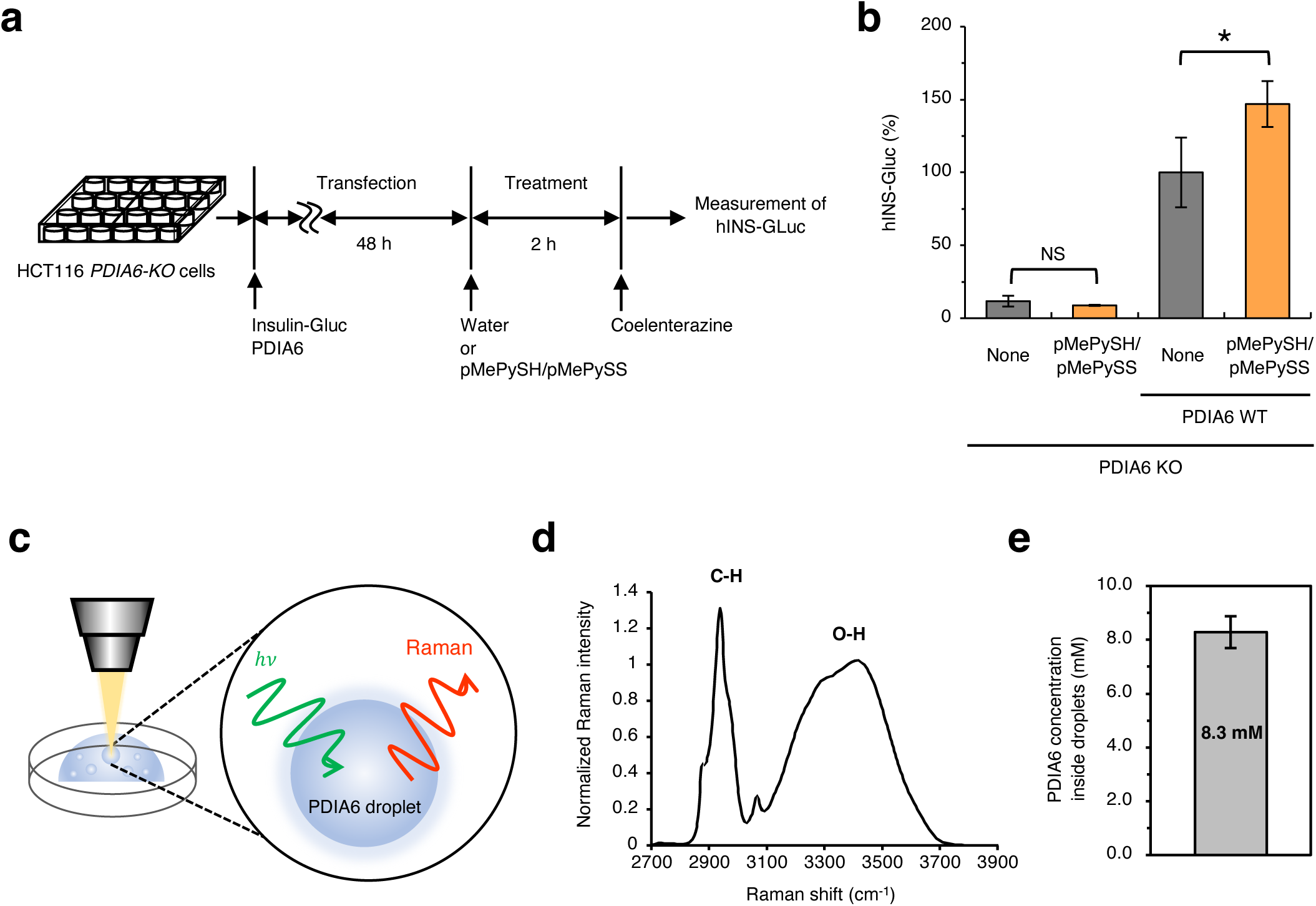
pMePySH/pMePySS increases insulin secretion by targeting PDIA6 enriched in foci. **a,** Scheme for measuring exogenous insulin secretion in *PDIA6*-KO cells in the presence or absence of PDIA6 following pMePySH/pMePySS treatment. **b,** Effects of GSH/GSSG and pMePySH/pMePySS on insulin secretion in HCT116 *PDIA6*-KO cells transfected with proinsulin-*Gaussia* luciferase (hINS-Gluc) with or without PDIA6. Cells were treated with 75 µM GSH/5 µM GSSG or 75 µM pMePySH/5 µM pMePySS for 2 h at 37°C. Data are presented as the mean ± s.d. from three independent experiments. Statistical significance was determined using a two-tailed unpaired Student’s *t*-test (*n* = 3). \**P*<0.05; NS, not significant. **c,** Direct Raman measurement of PDIA6 concentration within a single droplet. **d,** Averaged Raman spectrum of PDIA6 droplets. Data is presented as the mean of 16 particles from three independent experiments. **e,** PDIA6 concentration inside a droplet. Data are presented as the mean ± s.d. of 16 particles from three independent experiments.

## Discussion

The development of chemical tools ^32^ ^40^ ^41^, such as photoactivated protein oligomerization *in cellulo*, has enabled the precise manipulation of phase-separated droplets, revealing the biological significance of intracellular condensation ^42^. Beyond basic compartmentalization tools, recent advances have yielded peptide-, protein-, and chemical-based systems ^43^ that confer add-on functionalities to biomolecular condensates. For instance, photoresponsive liquid-solid phase transition ^44^, liquid-liquid phase separation ^45^, and macromolecular protein assembly ^46^ regulates the chemical reactivity of protein condensates under physiological conditions, demonstrating photoresponsive switching of condensate functionalities. While inhibitors of condensate formation can selectively attenuate enzymatic activity ^47^, specific chemical compounds—such as 4,4’-dianilino-1,1’-binaphthyl-5,5’-disulfonic acid, rosmanol quinone, and poly(4-styrenesulfonic acid-co-maleic acid) —can actively induce the formation of liquid-like droplets ^48^ ^49^ ^50^ and enhance enzymatic catalysis within them ^51^. As biomimetic compartmentalization agent, chloroplast-mimicking condensates enable highly efficient artificial photocatalytic water oxidation ^52^. Therefore, chemical strategies targeting phase-separated droplets represent a promising frontier for functional switching, regulating enzymatic networks, and establishing biomimetic platforms. Biomolecular condensates are commonly driven by weak, multivalent interactions between intrinsically disordered regions (IDRs), often classified as low complexity domains (LCDs). Fusion LCDs to enzymes ^53^, deploying LCDs alone ^54^ or utilizing multidomain scaffolding to drive liquid-like droplet formation ^55^ can drastically increase local enzyme concentration, thereby enhancing overall activity and facilitating the *de novo* design of synthetic enzymatic condensates. Enzymatic reactions within droplet substructures can be further manipulated by introducing specific amino acids to form internal vacuoles ^56^ or by engineering droplet-in-droplet architectures using IDR-tags ^57^, which underscores the dynamic, reaction-mediated rearrangement of liquid-like environments. Thus, LCDs and IDRs serve as highly versatile tags or fusion partners to endow artificial condensates with unique functionalities. Despite these major advances in functional compartmentalization tools, strategies capable of augmenting catalytic activity beyond inherent baseline levels within endogenous biological condensates have remained a long-standing challenge.

In this study, we established a chemical biology framework to target and boost the catalytic processing of reduced/denatured substrates within the phase-separated, liquid-like droplets of PDIA6, leveraging them as specialized reaction chambers for oxidative folding (Fig. 1). These condensates form in a Ca^2+^-dependent manner and maintain their structural integrity in the presence of standard GSH/GSSG redox couples without undergoing dissolution (Fig. 2c). Furthermore, the catalytic activity of PDIA6 droplets toward diverse client proteins—including proinsulin, BPTI, and antibodies—significantly outpaced that of dispersed PDIA6 under standard GSH/GSSG conditions, demonstrating that with condensation an optimized folding environment emerges (Fig. 2i). Importantly, chemically targeting the active-site CxxC motif of PDIA6 could amplify enzymatic efficiency within the droplet architecture. Regarding the development of enzymatic activators for dispersed PDIA6, we previously reported that pMePySH (p*K*a = 7.34, E°’ = -211 mV) enhances both the acidity and oxidizability of thiol groups, achieving a 2.6-fold acceleration in oxidative folding compared with glutathione ^35^. Remarkably, substituting GSH/GSSG with the pMePySH/pMePySS redox couple hyper-activated the internal machinery of PDIA6 droplets, yielding a 12-fold acceleration in folding kinetics and demonstrating a profound synergy between the chemical booster and the condensate microenvironment (Fig. 4g). Furthermore, pMePySH selectively targeted ER-localized PDIA6 foci *in cellulo* (Fig. 6), resulting in augmented oxidative folding efficiency and enhanced insulin secretion (Fig. 7b). This robust synergistic enhancement is connected to the phase-separated state of PDIA6. Raman spectroscopy showed that the internal PDIA6 concentration is enhanced to 400-fold (Fig. 7c–e and Extended Data Fig. 6a,b). The extraordinary increase in catalytic efficiency driven by a dual mechanism: the dramatic enrichment of the foldase within the droplet interior and the intrinsic hyper-activation of individual PDIA6 active sites by pMePySH/pMePySS.

Consequently, this work introduces a generalizable modality to manipulate protein folding pathways by targeting quality control granules with chemical compounds, offering critical insights into how functionalized small-molecule thiols can boost the performance of biological condensates.

Spatiotemporal control over protein quality control networks offers a powerful means to suppress the accumulation of cytotoxic, misfolded intermediates. Aberrant stagnation or spatial dysregulation of misfolded proteins frequently triggers pathological phase transitions, leading to the nucleation of amyloid fibrils or amorphous aggregates implicated in various protein misfolding diseases; artificially boosting the folding capacity within these quality control microenvironments represents a promising therapeutic avenue to intercept these pathological cascades. Furthermore, the oxidative folding platform established here holds immense potential for the biopharmaceutical industry, enabling the highly efficient production of disulfide-rich therapeutic agents, such as monoclonal antibodies, both *in vitro* and *in cellulo*. Continued exploration of such chemical biology approaches to drive high catalytic turnover within functional condensates will be essential to mitigate the structural risks associated with client accumulation and to advance the frontiers of synthetic and cell biology. Ultimately, this study establishes a paradigm for the chemical orchestration of biological condensates, demonstrating that tailored small molecules can interface with macromolecular phase separation to achieve non-linear gains in catalytic efficiency.

## Acknowledgements

We thank Prof. Kenji Inaba (Kyushu University) for PDIA6 plasmid, Prof. Taro Mannen (Ritsumeikan University) for providing U2OS cells stably expressing mCherry-PDIA6, Prof. Tsukasa Okiyoneda and Dr. Yuka Kamada (Kwansei Gakuin University) for supporting insulin secretion assay, Prof. Satoshi Ninagawa (Kobe University) for PDIA6 KO cell, Ms. Izumi Nagai (Tohoku University) for supporting the purification of PDIA6, and Ms. Kotone Ishii (Tohoku University) for supporting the purification of BPTI mutants. This research was funded by JSPS KAKENHI grants JP22H02205 (MO), JP23KK0105 (MO), JP21H05095 (MO), and JP21H05093 (TM); Japan Science and Technology Agency FOREST Program grants JPMJFR201F (MO) and JPMJFR2122 (TM); Japan Science and Technology Program for co-creating startup ecosystem grant JPMJSF2312 (MO); and grants from Takeda Science Foundation (MO); Mochida Memorial Foundation for Medical and Pharmaceutical Research (MO); Naito Foundation (MO); Uehara Memorial Foundation (MO); Terumo Life Science Foundation (MO); Astellas Foundation for Research on Metabolic Disorders (MO); Asahi Glass Foundation (MO); Mitsui Sumitomo Insurance Welfare Foundation (MO); Daiichi Sankyo Foundation of Life Science (MO); Sumitomo Foundation (MO); Ono Medical Research Foundation (MO); Nakatani Foundation (MO); and Shiraishi Foundation of Science Development (MO).

## Author Contributions Statement

Conceptualization: MO, Methodology: TM, and MO, Investigation: MW, TK, MF, NU, HA, KB, MH, KA, and TN, Data analysis: MK, Funding acquisition: TM and MO, Project administration: TM and MO, Supervision: TM and MO, Writing – original draft: MO, Writing – review & editing: MW, JB, TM and MO.

## Methods

### Recombinant protein expression and purification

A cDNA encoding human PDIA6 lacking the N-terminal signal sequence was subcloned into the pET15b vector between *Nde*I and *Bam*HI restriction enzyme sites (Novagen, Darmstadt, Germany). The plasmid incorporates an N-terminal 6-His tag. PDIA6 was overexpressed in *Escherichia coli* stain BL21 (DE3). Pelleted cells were resuspended in lysis buffer containing 50 mM Tris-HCl (pH 8.1) and 300 mM NaCl, and lysed by sonication. After centrifugation at 15,000*g* for 30 min at 4°C, the supernatant was applied to a Ni-NTA agarose column, and the target protein was eluted with lysis buffer supplemented with 200 mM imidazole. Eluted fractions were loaded onto a Resource Q column (Cytiva, Marlborough, MA, USA) for further purification. Finally, eluted fractions were purified by size-exclusion chromatography using a Superdex 200 increase column (Cytiva). Purified PDIA6 was confirmed by sodium dodecyl sulphate-polyacrylamide gel electrophoresis (SDS-PAGE). The PDIA6 solution was buffer-exchanged into 50 mM HEPES-NaOH (pH 7.2) and stored at -80°C until use. Wild-type BPTI (WT), the Cys/Ser-substituted BPTI mutant (BPTI_all-Cys/Ser_), and proinsulin were overexpressed in *E. coli* stain BL21 (DE3) and purified as previously described ^4 31 35^. DY-605-maleimide modification was performed following established protocols for ATTO-532-maleimide modification ^25^.

Expression of D3-L11 was performed as described previously ^37^. Briefly, the synthetic gene encoding D3-L11 was cloned into the pRA2 vector ^58^, and the protein was expressed in *E. coli* BL21 (DE3) using 0.5 mM IPTG induction. Cells were harvested by centrifugation at 7,000*g* for 15 min at 4°C. Cells were resuspended in buffer A (20 mM Tris-HCl (pH 8.0) and 500 mM NaCl) containing 5 mM imidazole, lysed by sonication, and centrifuged at 20,000*g* for 30 min at 4°C. The supernatant was filtered through a 0.2 µm filter and loaded onto a Ni-NTA agarose column equilibrated with buffer A. After washing with buffer A containing 100 mM imidazole, the target protein was eluted with buffer A containing 500 mM imidazole. The eluate was subjected to SEC on a HiLoad 26/600 Superdex 75 pg column (Cytiva) equilibrated with buffer A. The buffer was subsequently exchanged into 20 mM HEPES-NaOH (pH 7.0) and 50 mM NaCl via ultrafiltration.

### BPTI folding assay

Native BPTI (10 mg) was dissolved in 1 mL of 100 mM Tris-HCl (pH 8.0) containing 20 mM DTT and 8 M urea, and incubated for 3 h at 50°C. The reduced/denatured form of BPTI was purified by reversed-phase high-performance liquid chromatography (RP-HPLC) using an InertSustain C18 column (φ4.6×250 mm, GL science, Tokyo, Japan), and the collected fractions were lyophilized. The resulting powder was stored at -20°C until use. For oxidative BPTI folding by dispersed PDIA6, 20 µM PDIA6 was preincubated with 2 mM GSH and 0.1 mM GSSG in 50 mM HEPES-NaOH (pH 7.2) at 30°C. After 10 min, 5 µM of reduced/denatured form of BPTI was added, and reaction proceeded at 30°C for 0.5 to 60 min before being quenched with 0.5 M HCl. For oxidative BPTI folding within PDIA6 droplets, 20 µM PDIA6 was preincubated with 2 mM GSH and 0.1 mM GSSG in 50 mM HEPES-NaOH (pH 7.2) for 10 min at 30°C. Next, 3 mM CaCl_2_ was added to initiate phase separation, and the mixture was incubated for 10 min at 30°C. Reduced/denatured form of BPTI (5 µM) was then mixed with the PDIA6 droplets and incubated for 0.5 to 60 min at 30°C, followed by quenching with 0.5 M HCl. For assays utilizing low concentrations of thiol boosters, 20 µM PDIA6 was preincubated with 75 µM GSH or 75 µM pMePySH, along with 5 µM GSSG or 5 µM pMePySS, in 50 mM HEPES-NaOH (pH 7.2) for 10 min at 30°C. Droplet formation was incubated with 3 mM CaCl_2_ for 10 min at 30°C, and folding was initiated by adding 5 µM reduced/denatured BPTI. Reactions were quenched at designated time points (0.5 to 30 min) using 0.5 M HCl and 2 M urea. All folding assays were analyzed by RP-HPLC using a TSKgel Protein C4-300 column (Tosoh Bioscience), and absorbance was monitored at 229 nm. Peak identities were confirmed by matrix-assisted laser desorption/ionization time-of-flight mass spectrometry (MALDI-TOF/MS) analysis as previously described ^59^ _60._

### Substrate uptake and *in vitro* phase separation assays

PDIA6 droplet formation was initiated by mixing 50 µM PDIA6 and 4 mM CaCl_2_ in 50 mM HEPES-NaOH (pH 7.2). After 1 min, 1−10 µM BPTI WT or BPTI_all-Cys/Ser mutant_ was added to PDIA6 droplets for 5 to 60 min. For fluorescence visualization, 5 µM DY-605-labeled BPTI was added to the preformed droplets. Samples were placed on a Tomodish (Tomocube Inc., Daejeon, Korea) and monitored via holographic microscopy equipped with a HT-2H fluorescence system (Tomocube Inc.). Refractive index (RI) images represent the XY optical section of the central region of the droplets. The mean RI within individual droplets was quantified by averaging values across a circular region of interest (ROI) using TomoStudio (versions 3.2.8 and 3.3.9; Tomocube Inc.). To evaluate droplet behavior in the presence of foldable denatured substrates, 20 µM PDIA6 was preincubated with 2 mM GSH and 0.1 mM GSSG for 10 min. Reduced/denatured BPTI (5 µM) was then added, and droplets were imaged on a glass-bottom dish (Matsunami Glass Ind., Ltd., Osaka, Japan) using a CKX53 microscope (OLYMPUS, Tokyo, Japan).

To assess the effects of redox couples (GSH/GSSG vs pMePySH/pMePySS) on droplet formation, performed PDIA6 droplet (20 µM PDIA6, 3 mM CaCl_2_) were treated with 75 μM thiol (GSH or pMePySH) and 5 μM disulfides (GSSG or pMePySS) for 10 to 60 min, and examined via phase-contrast microscopy.

### Proinsulin folding assay

PDIA6 (20 μM) was preincubated for 10 min at 30°C in 50 mM HEPES-NaOH (pH 7.2) with or without 75 μM thiol (GSH or pMePySH) and 5 μM disulfide (GSSG or pMePySS). Droplet formation was initiated by adding 3 mM CaCl_2_. After 10 min, reduced/denatured proinsulin (5 μM) was mixed with PDIA6 droplets at 30°C. The oxidative folding reaction was quenched at specific time points by adding 2-aminoethyl methanethiosulfonate (7.0 mg/mL), a selective thiol functional group modification regent. Reaction mixtures were analyzed by RP-HPLC using a TSKgel ODS-100V 5 μm column (Tosoh Bioscience) with UV detection at 220 nm. Peak identities were verified by MALDI-TOF/MS as previously described ^59^ ^60^.

### Cell culture and apoptotic assays

U2OS/TR_mCherry-PDIA6 cells were cultured at 37°C under 5% CO_2_ in Dulbecco’s Modified Eagle Medium (DMEM) supplemented with 10% fetal bovine serum (FBS) and 1% penicillin/streptomycin^4^. To induce the expression of mCherry-PDIA6, the medium was supplemented with 15 ng/mL of doxycycline (Wako, Tokyo, Japan) for 24 h. For cytotoxicity evaluations, cells were seed onto 8-well chamber slides (ibidi, Gräfelfing, Germany), induced with doxycycline, and treated with either the GSH/GSSG couple (75 μM/5 μM), the pMePySH/pMePySS couple (75 μM/5 μM), or 5 mM H_2_O_2_ (as a positive apoptotic control) for 8 h at 37°C. Cells washed with PBS and incubated in FluoroBrite DMEM (Thermo Fisher Scientific) containing 4 mM L-glutamine (Nacalai Tesque, Kyoto, Japan), 10% FBS, CellEvent Caspase-3/7 Green ReadyProbes Reagent (Thermo Fisher Scientific) and 1 μg/mL Hoechst 333424 for 30 min. Confocal microscopic imaging was performed using an LSM 800 (Carl Zeiss AG, Oberkochen, Germany).

### Quantification of insulin secretion in HCT116 cells

Exogenous insulin secretion was quantified using a proinsulin-*Gaussia* luciferase (Gluc) reporter system ^61^. Subconfluent *PDIA6* knockout (KO) HCT116 cells in 24-well plates were co-transfected with pJNC-hINS-Gluc (RDB19845, obtained from RIKEN DNA Bank) and pcDNA3.1 encoding wild-type (WT) PDIA6 using Polyethylenimine (PEI) Max (Polysciences Inc., Warrington, PA, USA) for 48 h. Cells were then treated with 75 µM pMePySH and 5 µM pMePySS in culture medium for 2 h at 37°C. The medium was harvested and cleared of debris by centrifugation at 780*g* for 2 min. Secreted Insulin-Gluc activity was determined by mixing a 10 µL aliquot of the supernatant with 50 µL of PBS containing 5 ng/µL coelenterazine (CAS 55779-48-1, Santa Cruz Biotechnology, #sc-205904) in white 96-well plates. Luminescence was integrated for 0.7 s using a Luminoskan microplate reader equipped with an automated dispenser (Luminoskan; Thermo Fischer Scientifics, Waltham, MA, USA). To normalize secretion against total cellular protein, cells were lysed in RIPA buffer containing 1 mM phenylmethylsulfonyl fluoride (FUJIFILM Wako Pure Chemical Corporation), 5 µg/mL leupeptin (FUJIFILM Wako Pure Chemical Corporation), and 5 µg/mL pepstatin A (Peptide Institute, Inc.). Lysates were cleared at 14200*g* for 5 min, and protein concentrations were determined via standard colorimetric assays.

### Raman microscopic measurements and concentration determination

Raman spectra were acquired using a custom-built confocal Raman microscope. ^62^ ^63^ A frequency-doubled Nd:YVO_4_ CW laser (λ = 532 nm; Millennia Vs, Spectra-Physics, Santa Clara, CA, USA) served as the excitation source, coupled to an inverted microscope (IX-73; Olympus, Tokyo, Japan) and a thermoelectrically cooled CCD detector (DU416A-LDC-DDM; Andor, Belfast, UK). A 60 µL of droplet-containing sample was placed on a glass-bottom dish (Matsunami Glass, Osaka, Japan) pre-coated with poly[2-methacryloyloxyethyl phosphorylcholine-co-n-butyl methacrylate] to maintain droplet sphericity. To prevent evaporation, a reservoir of Milli-Q water was placed along the inner perimeter, and the chamber was sealed with a lid. Laser light was focused through a 100× oil-immersion objective lens (NA = 1.49, oil immersion, Olympus, Tokyo, Japan), which also collected the backscattered Raman signal. Signals were dispersed using an MS3504i polychromator (SOL instruments, Minsk, Belarus). The measuremens were performed as the confocal condition using the introduced pinholes. Exposure times inside and outside the droplets were set to 10 s per point with no accumulation, maintaining a laser power of 50 mW at the objective back- aperture.

To absolute-quantify the local macromolecular concentration within the droplets, the O-H stretching Raman band of water outside the condensates was utilized as an internal standard ^62^ ^63^. A calibration curve was constructed by measuring a concentration series of dispersed PDIA6 in HEPES buffer. The C-H stretching band of PDIA6 was integrated over the 2930-2960 cm^−1^ region and normalized to the O-H stretching band of water integrated over the 3350-3450 cm^-1^ region. Linear regression of the normalized intensities versus known PDIA6 concentrations yielded the calibration function. Multiple distinct spatial coordinates inside and outside the droplets were sampled. The partition concentration of PDIA6 within the condensates was determined by interpolating the normalized droplet C-H intensities onto the calibration curve, with the baseline contribution of the buffer component (HEPES) subtracted as the *y*-intercept.

---

**Extended Data Fig. 1.**
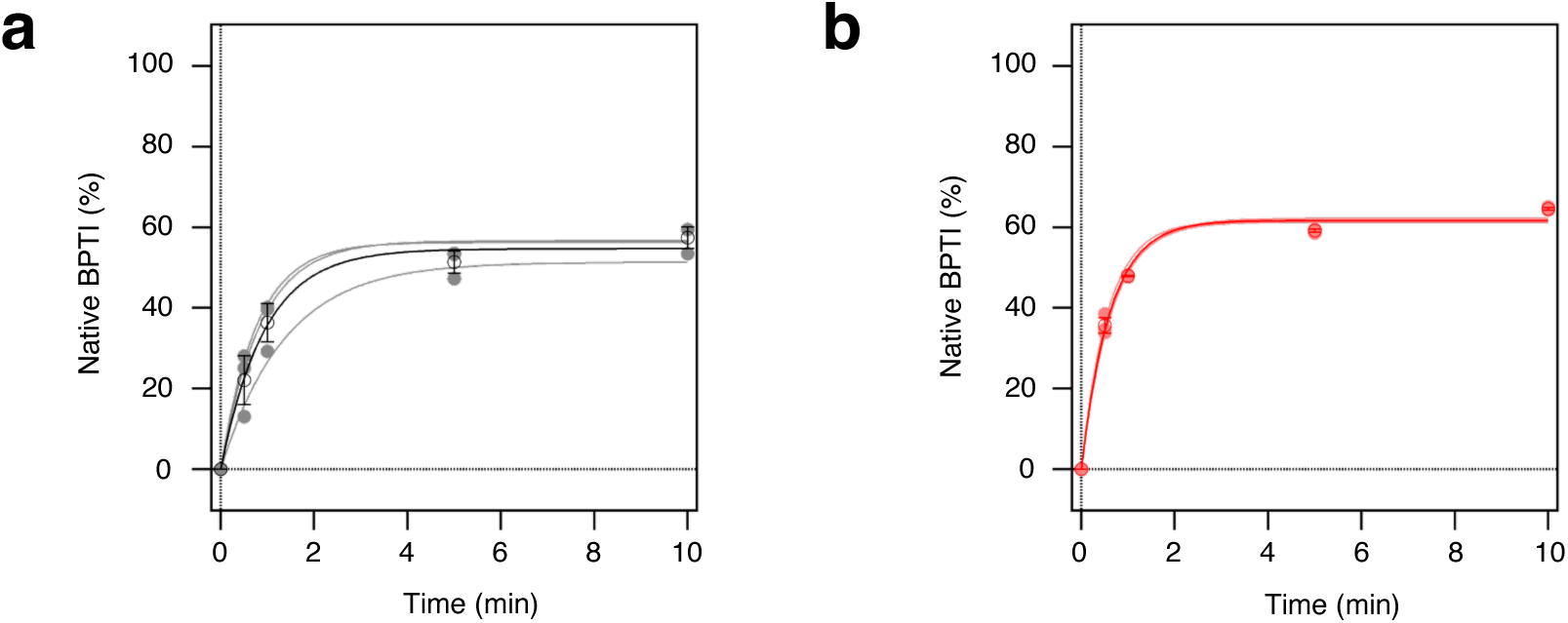
Kinetics analysis of oxidative BPTI folding catalyzed by dispersed or phase-separated PDIA6. **a,b**, Oxidative folding of BPTI by PDIA6 in the absence (**a**) or presence (**b**) of Ca^2+^ ions. Solid lines represent fitted curves. Rate constants were determined from three independent experiments (mean ± s.d). Rate constants were calculated by a single exponential curve fitting using IGOR Pro6 software (WaveMetrics, Inc., Lake Oswego, OR, USA, https://www.wavemetrics.com/).

**Extended Data Fig. 2.**
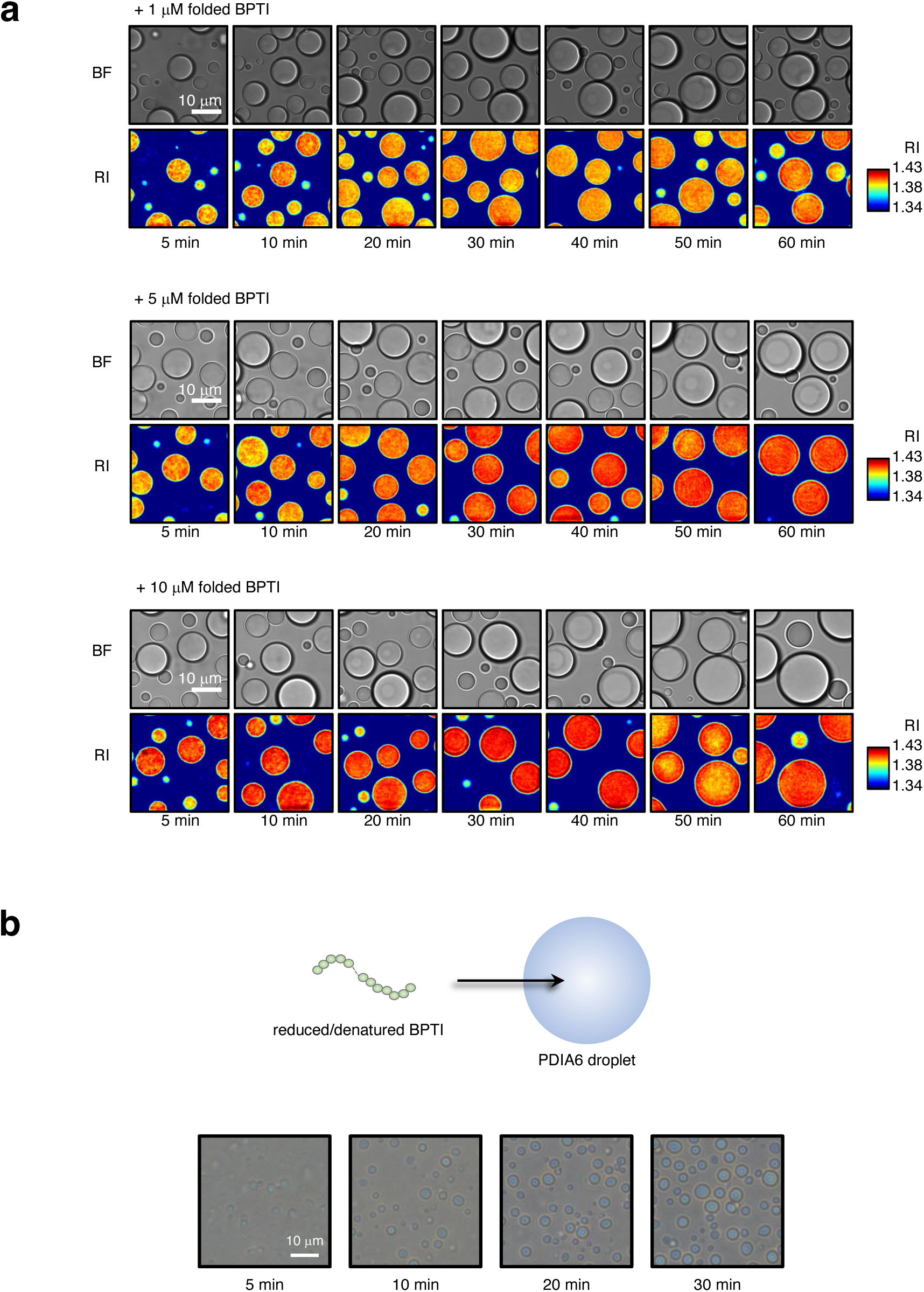
Uptake into PDIA6 condensates for folded and reduced/denatured BPTI. **a,** Bright field (BF) and refractive index (RI) images of PDIA6 condensates in the presence of native BPTI. Native BPTI (1−10 µM) was added to PDIA6 droplet solution containing 50 µM PDIA6 and 4 mM CaCl_2_ at pH 7.2. **b,** Time-course images of PDIA6 droplets in the presence of reduced/denatured BPTI. Reduced/denatured BPTI (5 µM) was added to PDIA6 droplet solution containing 20 µM PDIA6, 3 mM CaCl_2_, 2 mM GSH and 0.1 mM GSSG at pH 7.2. **a,b,** These experiments were independently repeated at least three times.

**Extended Data Fig. 3.**
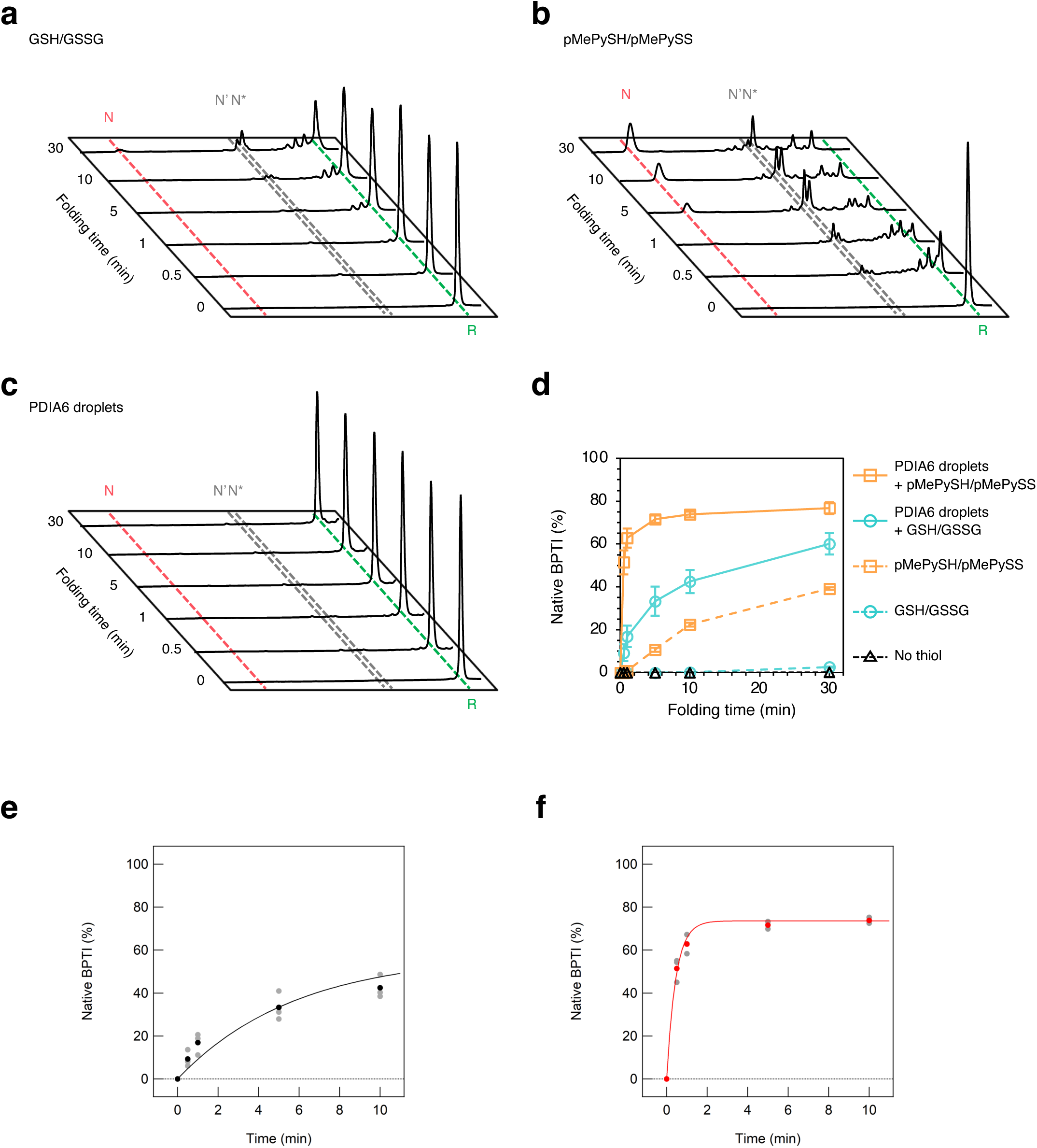
Kinetics analyses of oxidative BPTI folding via catalysis by thiol/disulfide compounds and PDIA6 droplets. **a−c**, Time-course reversed-phase HPLC analyses of oxidative BPTI folding by GSH/GSSG (**a**), pMePySH/pMePySS (**b**) and PDIA6 droplets (**c**). N, N’, N*, and R represent native BPTI, folding intermediates of BPTI with two disulfide pairings (30–51 and 14–38 for N’, and 5–55 and 14–38 for N*), and reduced/denatured forms of BPTI, respectively. These experiments were performed three times independently. **d,** Time-course plots of the formation of native BPTI via catalysis by GSH/GSSG, pMePySH/pMePySS, PDIA6 droplets, PDIA6 droplets with GSH/GSSG, and PDIA6 droplets with pMePySH/pMePySS. Data are presented as the mean ± s.d of three independent experiments. **e,f**, Time-course plots of the formation of native BPTI via catalysis by PDIA6 droplets in the presence of GSH/GSSG (**e**) or pMePySH/pMePySS (**f**). Solid lines represent fitted curves. Gray circles represent individual data points, and black and red circles represent average data (e and f, respectively). Rate constants were determined from three independent experiments (mean ± s.d). Rate constants were calculated by single exponential curve fitting using IGOR Pro6 software (WaveMetrics, Inc.).

**Extended Data Fig. 4.**
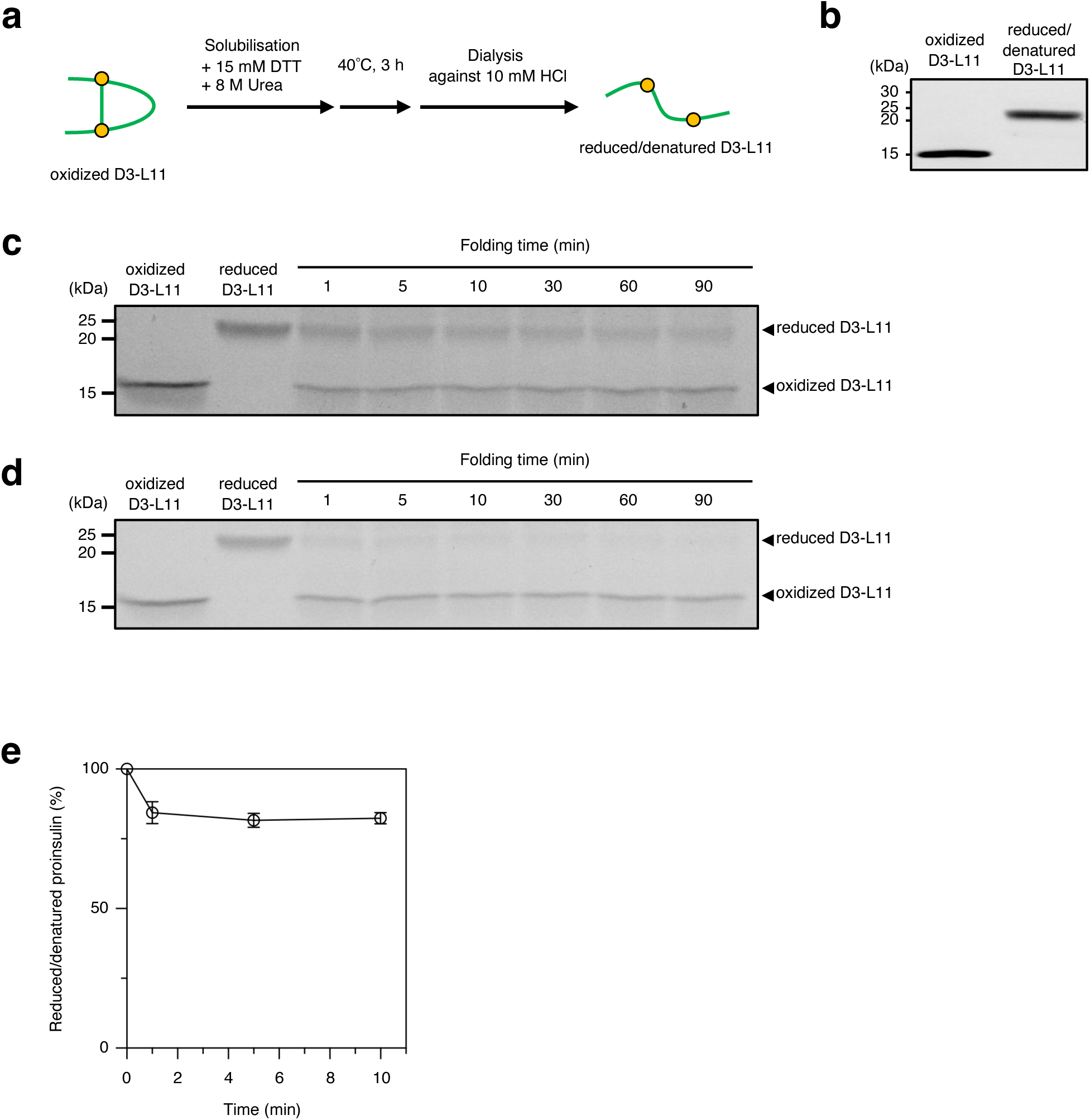
Oxidative folding of various substrates by PDIA6 droplets with pMePySH/pMePySS. **a,** Preparation of reduced and denatured D3-L11. **b,** Redox states of D3-L11 pre- and post-solubilisation and dialysis. Oxidized and reduced/denatured D3-L11 were modified with 2 kDa maleimide-PEG. Redox states were analysed by non-reducing SDS-PAGE. **c,d,** Time course of D3-L11 redox states during oxidation catalysed by PDIA6 droplets in the presence of GSH/GSSG (**c**) or pMePySH/pMePySS (**d**). Reduced/denatured D3-L11 (0.7 µM) was mixed with PDIA6 droplet solution containing 20 µM PDIA6, 3 mM CaCl_2_, and either 50 µM GSH/ 20 µM GSSG or 50 µM pMePySH/ 20 µM pMePySS at pH 7.2. **e**, Time-course plots of reduced/denatured proinsulin catalysed by PDIA6 droplets without thiol/disulfide compounds. Data are presented as the mean ± s.d of three independent experiments. **b–e,** These experiments were performed three times independently.

**Extended Data Fig. 5.**
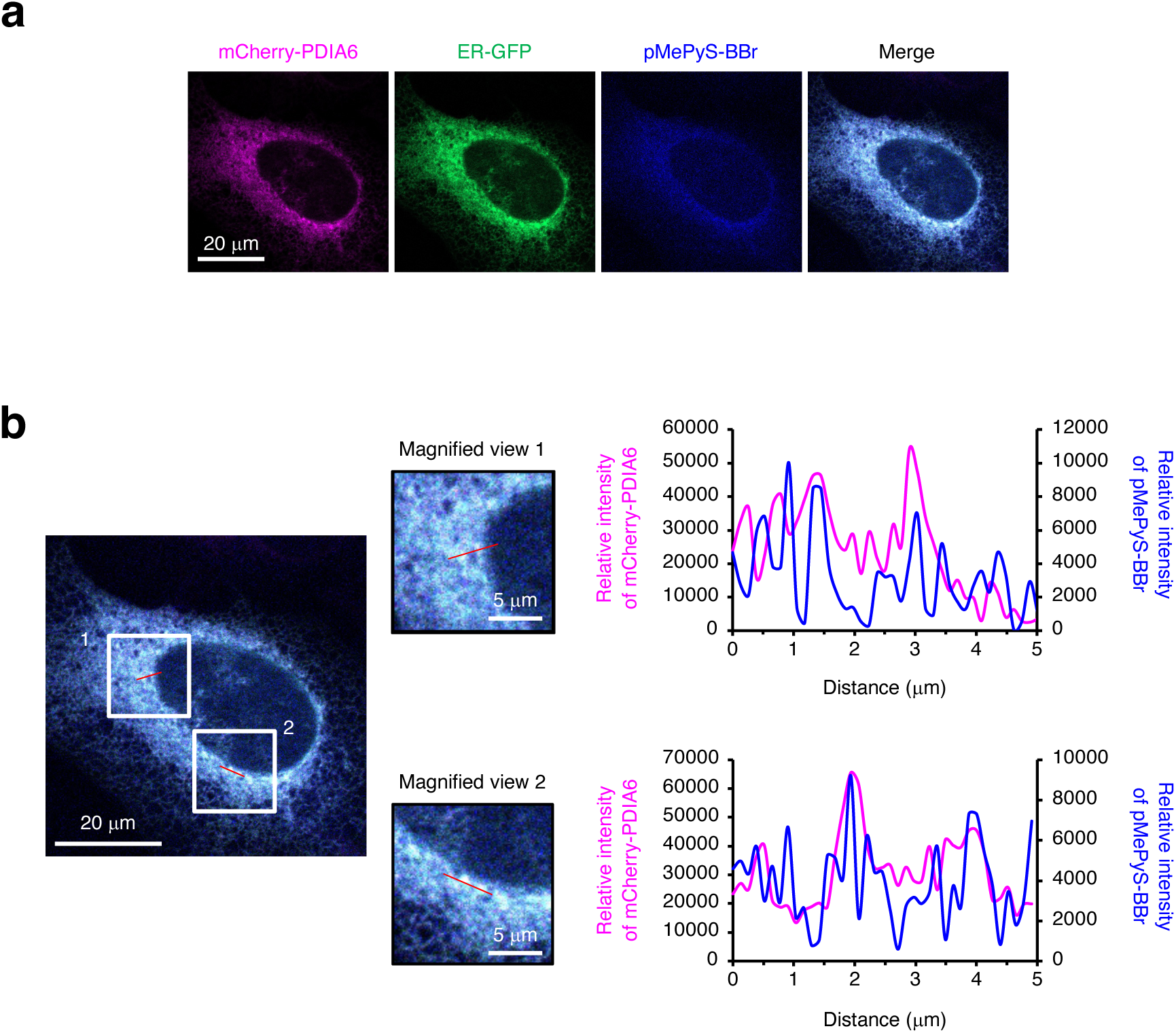
Co-localization of PDIA6 and pMePySH in the ER. **a,** U2OS cells stably expressing mCherryPDIA6 were stained with CellLight ER-GFP and were treated with 75 µM pMePyS-BBr for 5 h. **b**, Line scan analyses along the red lines indicated in the merged images (reproduced from **a**).

**Extended Data Fig. 6.**
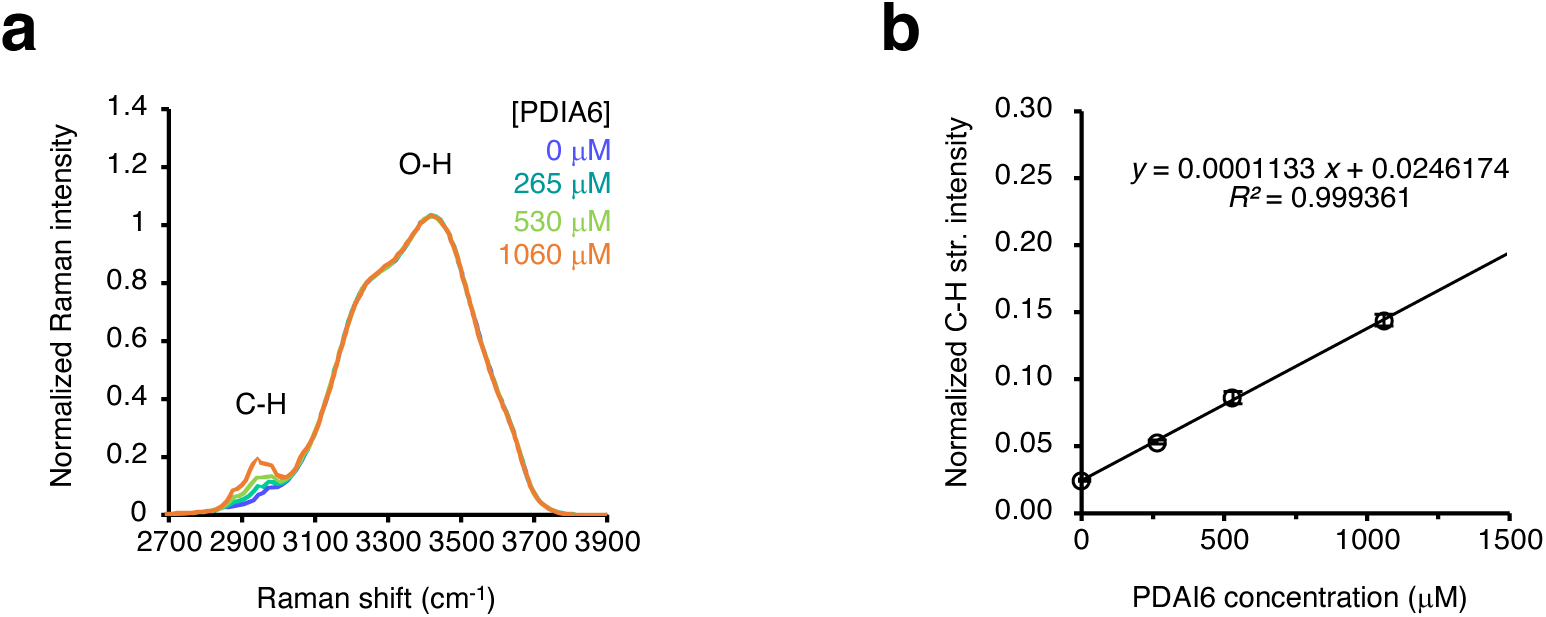
Raman spectra of dispersed PDIA6. **a,** Raman spectra of dispersed solutions of PDIA6 at different concentrations. **b**, A calibration line constructed by plotting the intensity of the C–H stretching band, normalized to the intensity of the O–H stretching band of water, against PDIA6 concentration. Data are presented as the mean ± s.d. of 15 measurements from three independent experiments.

