## Supplementary material for "Chemical Boosting of Foldase Condensates Accelerates Oxidative Protein Folding": Supp_Fig

### Extended Data Fig. 1

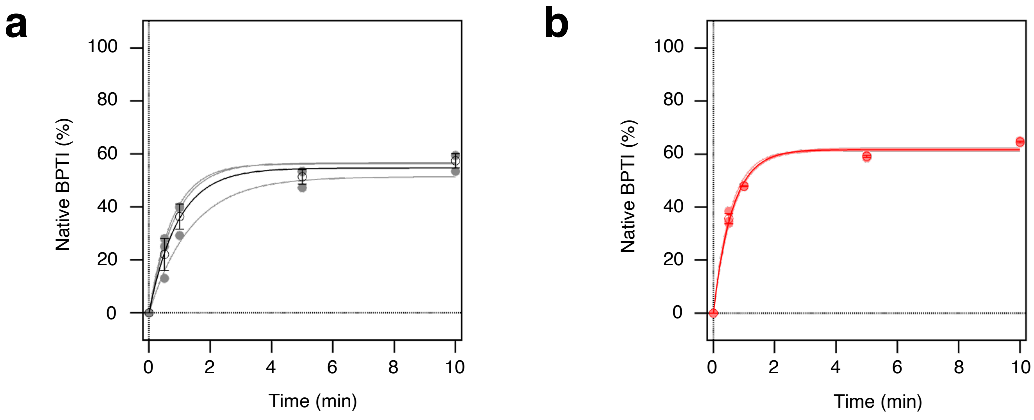

### Extended Data Fig. 2

**a**

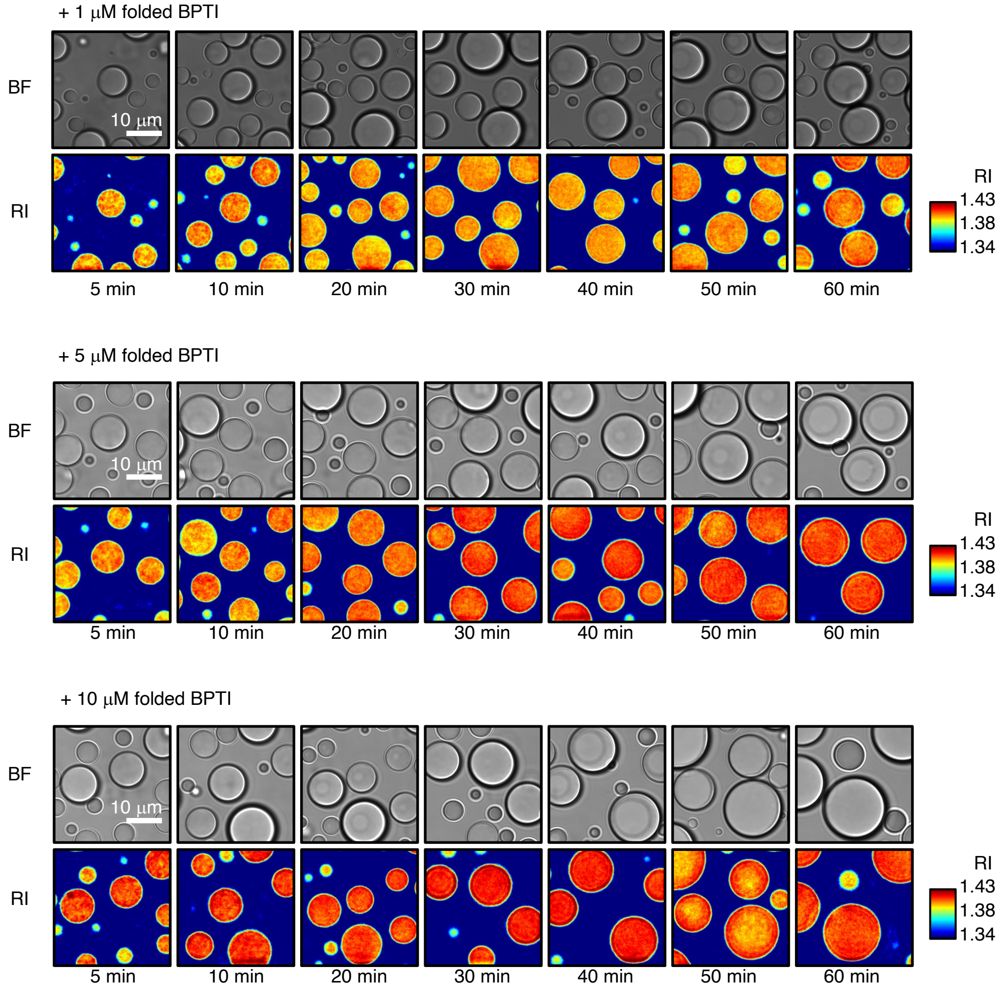


**b**

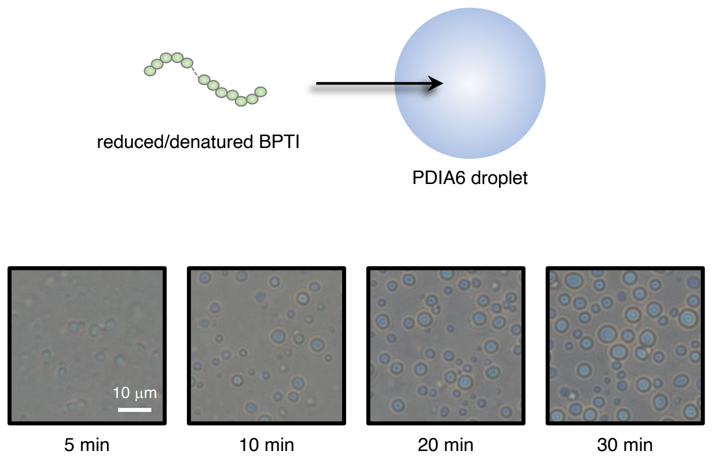

### Extended Data Fig. 3

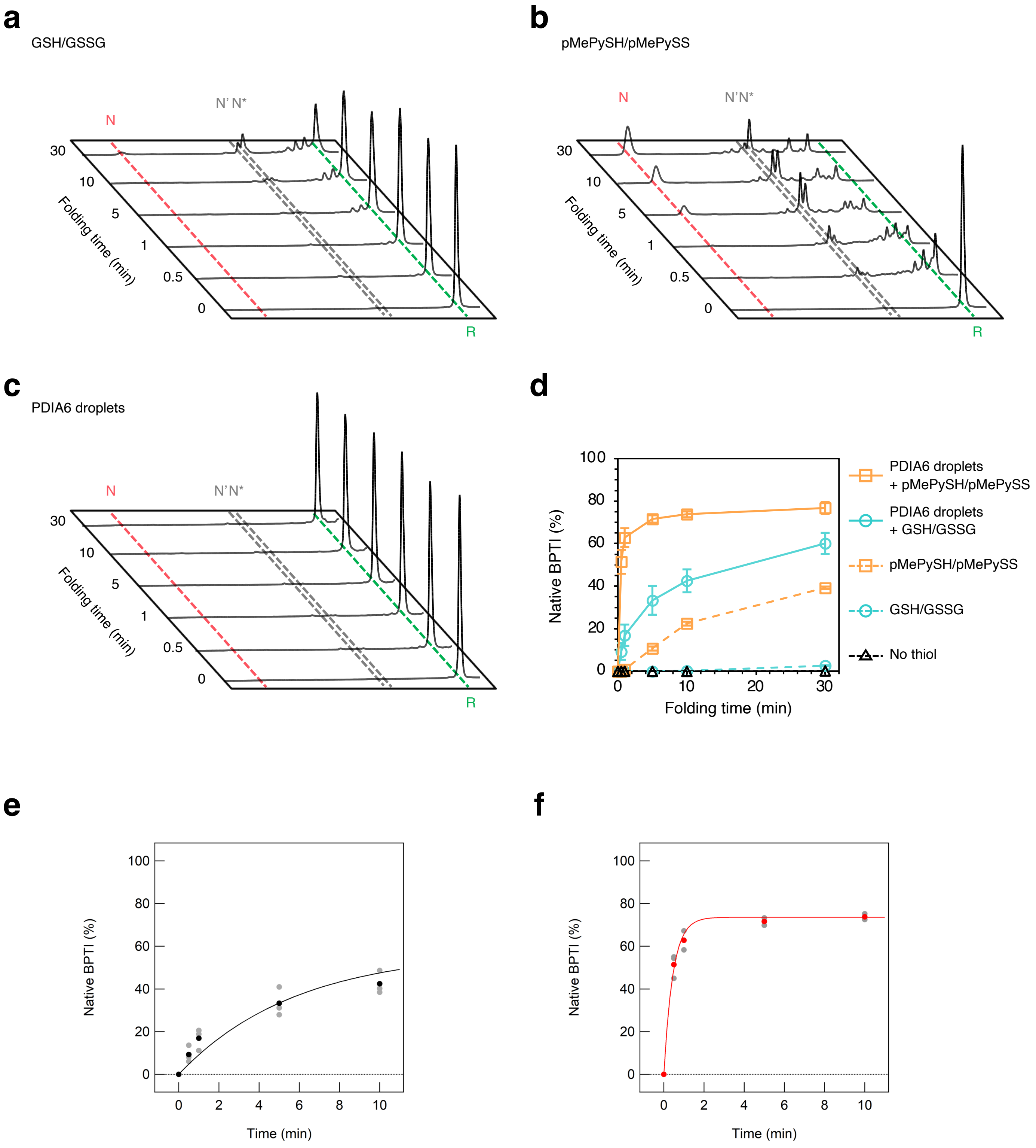

### Extended Data Fig. 4

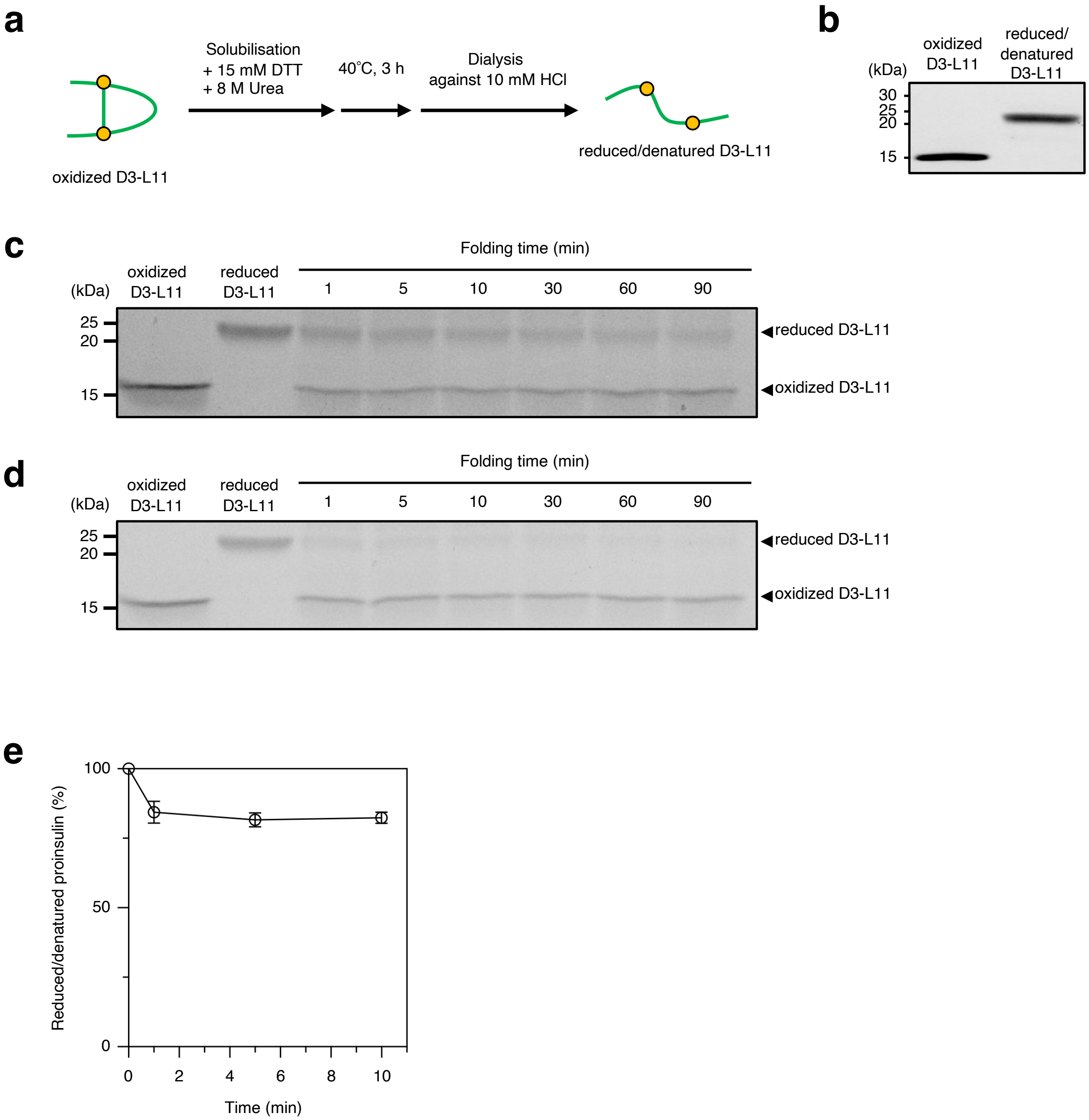

### Extended Data Fig. 5

**a**

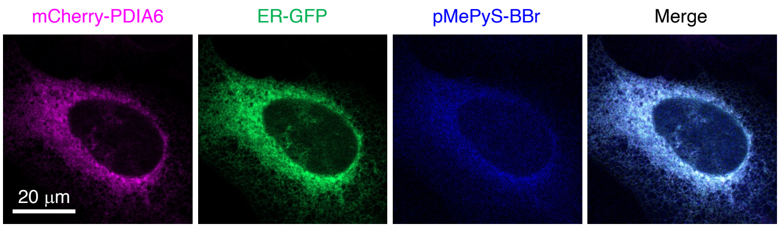


**b**

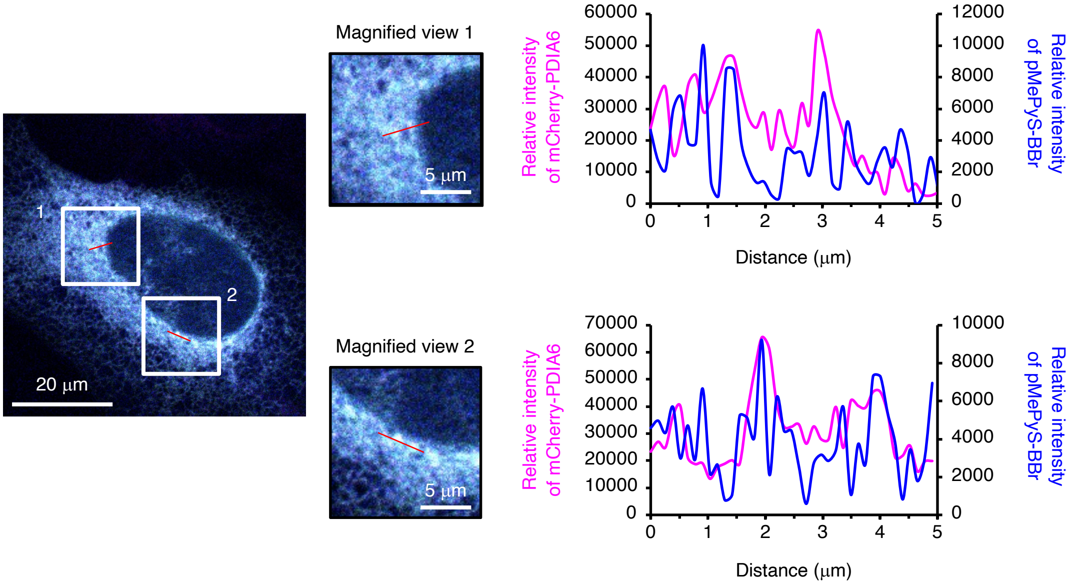

### Extended Data Fig. 6

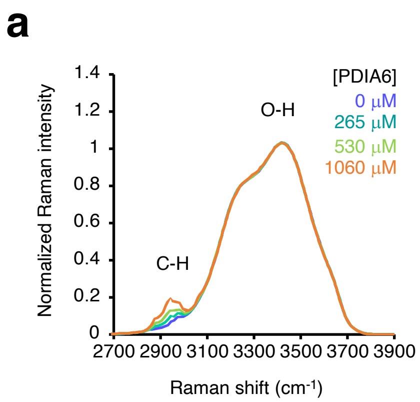


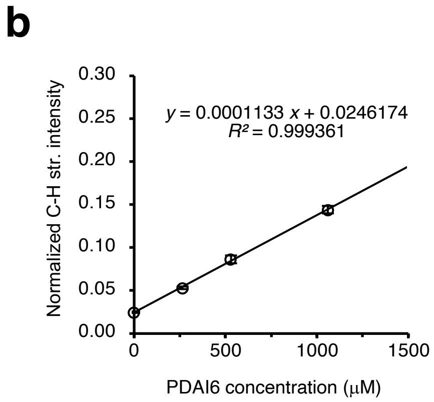
